# *In vitro* approaches to study centriole and cilium function in early mouse embryogenesis

**DOI:** 10.1101/2025.03.21.644416

**Authors:** Isabella Voelkl, Tamara Civetta, Mirijam Egg, Marie Huber, Songjie Feng, Alexander Dammermann, Christa Buecker

## Abstract

Although centrioles and primary cilia play an essential role in early mammalian development, their specific function during the interval between their initial formation and the subsequent arrest of embryogenesis in embryos deficient in centrioles or cilia remains largely unexplored. Here, we demonstrate that different 3D *in vitro* model systems recapitulate early centriole and cilium formation in mouse development. Centrioles and cilia are dispensable in 3D *in vitro* mouse rosettes, a model system that mimics key events of implantation, including polarization and lumenogenesis. In gastruloids, a model system that recapitulates developmental processes up to 8.5 days after fertilization, centriole loss results in early disassembly. In contrast, cells devoid of cilia continue to form elongated, differentiated and polarized gastruloids, with minor differences at 96 h. Finally, we show that in a mutant affecting the centriolar distal appendages, cilia are absent from 2D cultures but are capable of forming in 3D rosettes and gastruloids, highlighting the importance of multifactorial 3D environment setups in developmental studies.

**Summary:** This study presents the first *in vitro* analysis of centriole and cilium formation during early mouse embryonic development, using 3D models to mimic implantation, tissue patterning, and axis elongation, offering a controlled platform for investigating their roles in embryogenesis.

## Introduction

Centriole-based centrosomes serve as the predominant microtubule organizing centers (MTOCs) in animal cells, playing a vital role in mitotic spindle assembly and cell division (Bornens, 2012; Nigg & Raff, 2009). In addition, centrioles also provide the structural foundation of primary cilia - solitary, antenna-like sensory organelles that project from the surface of most vertebrate cells to mediate cellular signaling (Sorokin, 1968). Cilium formation requires the gradual maturation of centrioles over two cell cycles, enabling the mother centriole to transform into the basal body (Kong et al., 2014; Xiao et al., 2020). In mammals, this process depends on the distal appendage protein CEP83, which facilitates the docking of the basal body to the plasma membrane (Lo et al., 2019; Tanos et al., 2013). This connection then serves as a structural template for the extension of the axoneme, the microtubule core, with the assistance of the intraflagellar transport (IFT) machinery, including the IFT-B core component IFT88 (Haycraft et al., 2007). Although the ciliary membrane comprises only around 1/200^th^ of the total cell surface area, it is highly enriched in receptors, channels and effectors, essential for homeostasis and tissue patterning during embryogenesis. These components are crucial for mediating signal transduction, including pathways such as Hedgehog (Hh) and Wnt signaling, in particular during development (Mill et al., 2023). Consequently, dysfunction of primary cilia can give rise to pleiotropic and sometimes severe disorders, collectively termed ciliopathies, which can affect multiple organs, including kidneys, eyes, liver, brain, heart, lung and skeleton (Reiter & Leroux, 2017). To understand how centrioles and primary cilia develop and acquire their function during mouse embryonic development, it is essential to dissect their emergence within the embryo.

In contrast to human, *C. elegans* and zebrafish development, in which centrioles are indispensable (Avidor-Reiss et al., 2019; Pelletier et al., 2006; Yabe et al., 2007), the initial cell divisions of a developing mouse embryo after fertilization occur in the absence of centrioles, as both the oocyte and the sperm undergo centriole degeneration (Gueth-Hallonet et al., 1993; Manandhar et al., 1999). The first acentriolar foci of pericentriolar material (PCM) appear at the morula stage on embryonic day E2.5, and centrioles are formed *de novo* slightly later at the blastocyst stage at E3.5 (Howe & FitzHarris, 2013). Primary cilia emerge even later and are first detected on epiblast cells after implantation, around E5.5-E6, coinciding with cavitation and the onset of gastrulation. By E6, over 30% of epiblast cells are ciliated, and by E8, primary cilia are present on cells of all three germ layers, including the node (F. K. Bangs et al., 2015), the site where the left-right body axis is established (Brennan et al., 2002).

Centriole and cilium loss is lethal in mouse embryos. Homozygous *Plk4* KO mouse embryos lacking centrioles arrest in development at E7.5, accompanied by delays in cell division and ultimately apoptosis (Hudson et al., 2001). Mutations in IFT proteins inevitably result in mouse embryonic lethality by mid-gestation, around E11 (Murcia et al., 2000), due to the disruption of essential signaling pathways such as Wnt and Hh (Cortellino et al., 2009; Huangfu et al., 2003). Although primary cilia play essential roles in early mammalian development (Amack, 2022; F. Bangs & Anderson, 2017; Gerdes et al., 2009), their specific functions during the interval between their initial formation and the subsequent arrest of embryogenesis in cilia-deficient embryos remain largely unexplored. This limited understanding stems from the challenges associated with studying primary ciliogenesis *in vivo*, as these critical stages of development are difficult to observe *in situ* in the living embryo, offering only a limited, static perspective of developmental processes.

Mouse embryonic stem cells (mESCs) are a well-established *in vitro* model to study different aspects of mouse embryonic development. They are derived from the inner cell mass (ICM) of the embryo, molecularly resembling the pre-implantation epiblast (Nichols & Smith, 2009) and can be differentiated into all three germ layers. The first differentiation step is the transition into formative, epiblast-like cells (EpiLCs) (Buecker et al., 2014; Hayashi et al., 2011). mESCs serve as a powerful tool to generate various 3D *in vitro* model systems, including the embedded rosette model, which mimics critical aspects of implantation such as polarization and lumenogenesis (Bedzhov & Zernicka-Goetz, 2014; Shahbazi et al., 2017), and gastruloids, a developmental model system which recapitulates early embryonic cell fate decisions up to approximately 8.5 days post-fertilization (Beccari, Moris, et al., 2018).

In this study, we used 3D *in vitro* rosettes and mouse gastruloids as model systems to investigate the formation and function of centrioles and cilia during early mouse embryonic development. We show that differentiation *per se* is not a driver of cilium formation during transition of mESCs to EpiLCs; only in combination with polarization and lumenogenesis is ciliogenesis enhanced. Cilium- (*Ift88* KO, *Cep83* KO) and centriole (*Plk4* KO)-deficient rosettes maintain rosette-like morphology with a central lumen. However, *Plk4* is indispensable for gastruloid formation, as its loss leads to gradual disassembly in homozygous mutants. In contrast, *Cep83* and *Ift88*-deficient mESCs develop into elongated gastruloids, with only minor morphological differences at 96 h. Finally, a CEP83 truncation mutant (*CEP83Δexon4*) reveals surprising differential phenotypes between 2D cultures and 3D gastruloids, highlighting the importance of 3D models in developmental studies. This study provides the first comprehensive analysis of primary ciliogenesis and centriole formation employing *in vitro* differentiation models of early development.

## Results

### Primary ciliogenesis during exit from naive pluripotency

In the mouse embryo, cilia first arise on epiblast cells after implantation at the time of cavitation at E5.5-E6. In contrast, mESCs already possess the capacity to form primary cilia, which are found in a small proportion (∼5%) of cells (Bangs et al., 2015); however, it remains unclear whether cell differentiation enhances ciliogenesis *in vitro*. Removal of 2iLIF irreversibly differentiates naive mESCs into epiblast-like cells (EpiLCs) (Buecker et al., 2014; Hayashi et al., 2011), a process described as transitioning into the formative state of pluripotency or the exit from naive pluripotency (Smith, 2017). Here, we tested the potential of primary cilia to form during the transition of naive mESCs to formative EpiLCs (Fig. 1 A). We differentiated mESCs into EpiLCs for 48 h by removal of 2iLIF and determined their ciliation rate based on immunofluorescence staining using antibodies against the ciliary marker ARL13B and acetylated α-tubulin, a marker for stable tubulin (Fig. 1 B). Under mESC conditions, approximately 5% of cells exhibited a cilium, in line with previous reports. This ciliation rate remained unchanged during the differentiation into EpiLCs, indicating that the transition into formative pluripotency itself is not a driver of ciliogenesis (Fig. 1 C). The length of primary cilia of both mESCs and EpiLCs was between 1 and 2 µm, with EpiLCs exhibiting moderately shorter cilia. In summary, the differentiation of mESCs towards EpiLCs does not increase ciliogenesis.

**Fig 1.**
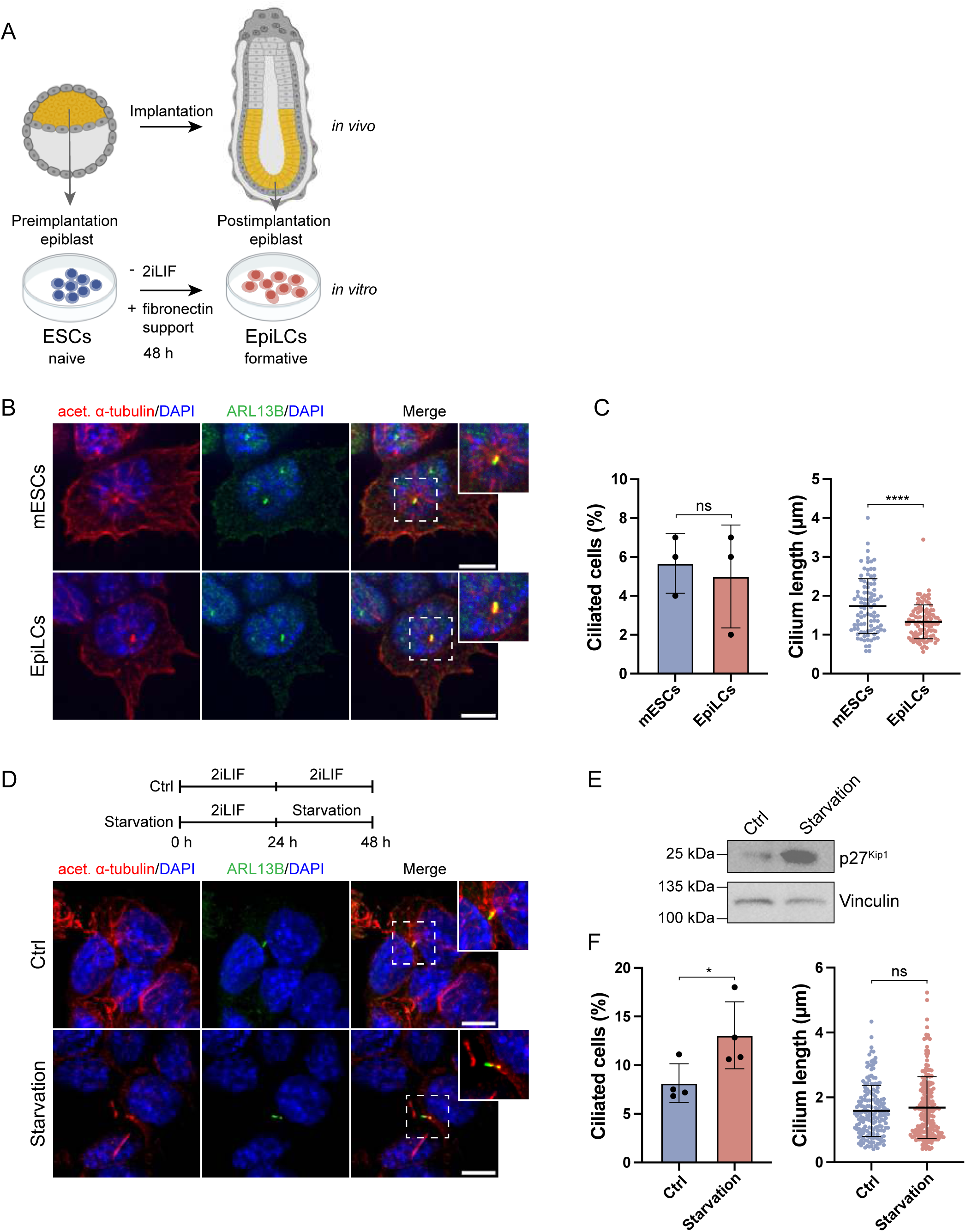
Exit of naive pluripotency does not increase primary ciliogenesis. (A) Schematic overview of ESC to EpiLC transition. Upper panel depicts the corresponding cell fate decisions *in vivo*, lower panel the conditions for the exit of the naive state to formative EpiLCs *in vitro*. (B) Representative examples of immunofluorescence staining of naive mESCs and formative EpiLCs after 48 h of differentiation labeled with antibodies against acetylated α-tubulin (red), the ciliary marker ARL13B (green) and DAPI staining (blue). Maximum intensity projection of central z-planes. (C) Ciliated cells (%) and cilium length (µm), mean + SD from n = 3 independent experiments (****p < 0.0001, unpaired t-test). (D) Representative examples of immunofluorescence staining of naive control mESCs (Ctrl) and after 24 h serum starvation, labeled with antibodies against acetylated α-tubulin (red), ARL13B (green) and DAPI staining (blue). Maximum intensity projection of central z-planes. (E) Western blot analysis of starvation marker p27 in naive control ESCs and after 24 h starvation. Vinculin was used as a loading control. (F) Ciliated cells (%) and cilium length (µm), mean + SD from n = 3 independent experiments (*p < 0.05, unpaired t test). Scale bar (B, D): 10 µm.

Ciliogenesis increases in response to serum starvation in various cell types such as retinal pigmented epithelium (RPE) cells and human embryonic kidney (HEK) cells (Takahashi et al., 2018). Since differentiation did not increase the rate of ciliogenesis, we investigated whether serum starvation could enhance ciliogenesis in our mESC model. We cultivated mESCs for 24 h and removed serum and additives (Methods, Cell starvation) for an additional 24 h before assessing cilium formation via immunofluorescence microscopy (Fig. 1 D). Starvation itself was monitored by the expression of p27 (Fig. 1 E), a cyclin-dependent kinase inhibitor that plays a crucial role in regulating the cell cycle and maintaining cellular quiescence. Elevated p27 levels are often associated with cell cycle arrest, particularly during starvation-induced quiescence (Li et al., 2019). Serum-deprived mESCs significantly increased ciliation compared to non-starved cells while cilium length remained unaffected (Fig. 1 F).

Together, these results suggest that a low percentage (5%) of naive mESCs is capable of forming cilia. The efficiency of ciliogenesis is not elevated during cell differentiation of mESCs to EpiLCs but can be increased by starvation, similar to observations in other cell types.

### 3D *in vitro* rosettes as a model system for primary ciliogenesis

*In vivo*, cilia first arise on embryonic epiblast cells after implantation at the time of cavitation at E5.5-E6 (F. K. Bangs et al., 2015). During these developmental steps, the cells undergo transcriptional and morphological changes, such as polarization. While EpiLCs are an excellent model system to study the transcriptional changes associated with exit from naive pluripotency, they do not recapitulate the morphological transformations occurring *in vivo*. Implantation itself is not tractable *in vivo*; therefore, we set out to find a suitable *in vitro* model to study early mammalian ciliogenesis in a controlled and monitored environment. We adapted the previously published 3D *in vitro* rosette assay (Bedzhov & Zernicka-Goetz, 2014; Shahbazi et al., 2017), a model for polarization and lumen formation, as a system to study ciliogenesis in early mouse embryonic development. In the 3D *in vitro* rosette assay, mESCs were embedded in Basement Membrane Extract (BME) without 2iLIF for up to 72 h to induce cell differentiation and lumenogenesis (Fig. 2 A). Under 2iLIF conditions, lumenogenesis was inhibited in 3D rosettes, as evidenced by the absence of the luminal marker PODXL. In contrast, removal of 2iLIF promoted rosette-shaped organization of cells and lumenogenesis (Fig. 2 B). We quantified the timing of lumenogenesis and evaluated the efficiency of rosette formation under differentiation conditions to assess the robustness of this model system. After 48 h, 56% of rosettes exhibited lumen formation, increasing to 100% by 72 h (Fig. 2 C), demonstrating the system’s high reproducibility and efficiency. Investigating the suitability of this system to study ciliogenesis, we immunostained rosettes after 72 h with antibodies against ARL13B and acetylated α-tubulin (Fig. 2 D). Both markers revealed prominent rod-shaped primary cilia, protruding into the lumen of the rosettes.

**Fig 2.**
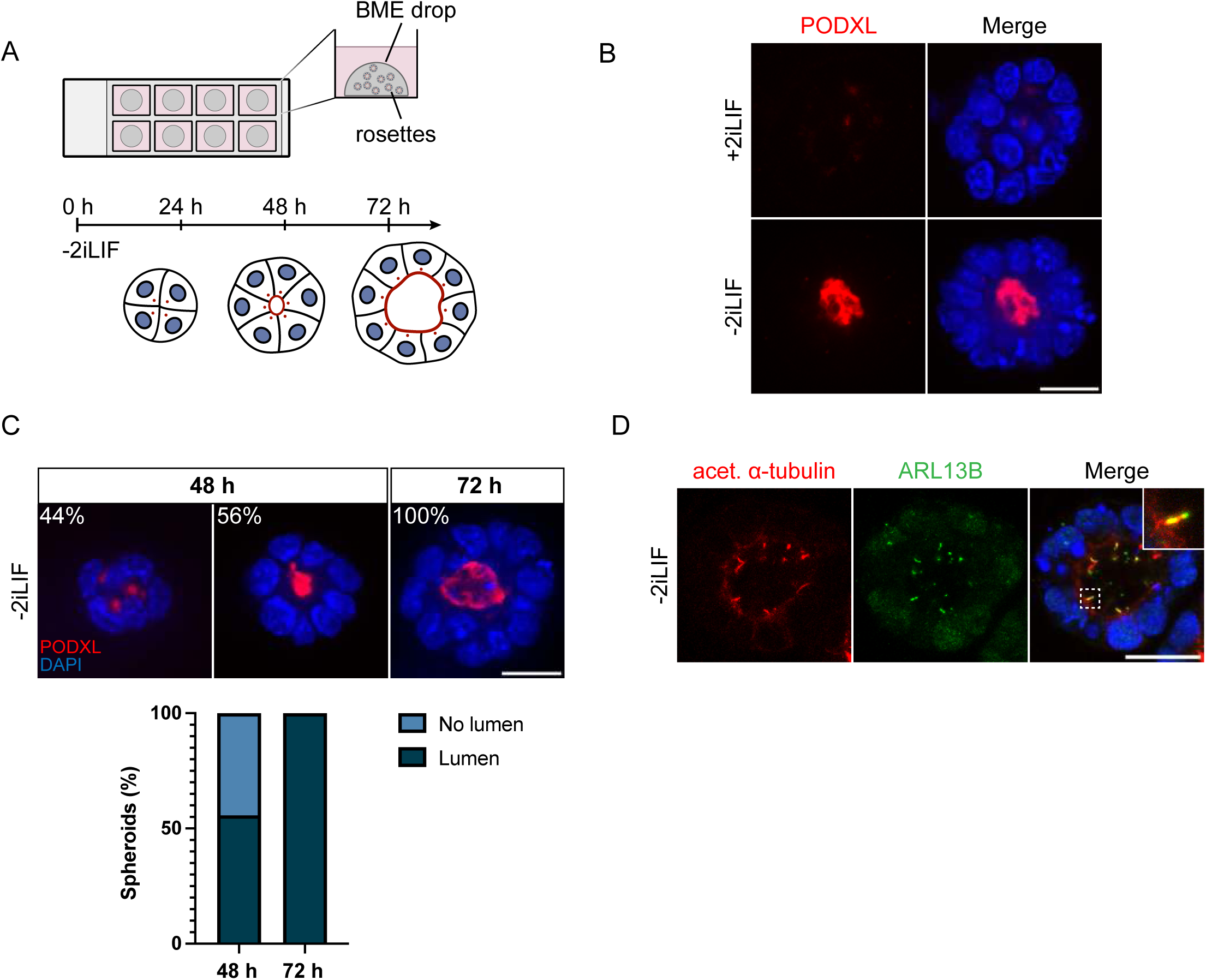
3D rosette assay recapitulates lumen and cilium formation *in vitro*. (A) Outline of the 3D *in vitro* mESC rosette assay, based on (Bedzhov & Zernicka-Goetz, 2014; Shahbazi et al., 2017) (B) Immunofluorescence staining of BME-embedded 3D *in vitro* rosettes growing under +/- 2iLIF conditions for up to 72 h. Representative example of immunofluorescence staining of rosettes, labeled with antibodies against lumen marker PODXL (red) and DAPI staining (blue). The central z-plane is depicted. (C) Representative immunofluorescence staining and timing of lumenogenesis in mESC-derived rosettes without 2iLIF at 48 h and 72 h. Antibodies against PODXL (red) and DAPI staining (blue) are indicated. Lumen quantification (%) from n=2 independent experiments. The central z-plane is depicted. (D) Immunofluorescence staining of rosettes growing without 2iLIF for 72 h, labeled with antibodies against acetylated α-tubulin (red), ARL13B (green) and DAPI staining (blue). Maximum intensity projection of central z-planes. Scale bar (B, C, D): 20 µm.

In summary, the 3D *in vitro* rosette system is ideally suited for investigating early ciliogenesis during mouse embryonic development *in vitro*, integrating key developmental processes such as cell polarization, lumenogenesis, and the dynamics of cilium assembly.

### Depletion of cilia and centrioles in mESCs to investigate early centriolar/ciliary functions

Next, we sought to investigate the role of centrioles and cilia in early mouse embryonic development. Using CRISPR/Cas9, we deleted the ciliary protein Intraflagellar Transport 88 (IFT88) (Fig. 3 A), a component of the intraflagellar transport machinery essential for cilium assembly and function (Pazour et al., 2000). Additionally, we targeted centrioles by deleting Polo-Like Kinase 4 (PLK4), a serine/threonine kinase with a critical role in regulating centriole duplication (Bettencourt-Dias et al., 2005; Habedanck et al., 2005; Swallow et al., 2005). In our KO approach, we designed two gRNAs for the *Ift88* KO, targeting exon 7 and the following intron, and two gRNAs for deletion of *Plk4*, specific to exon 5 and the following intron (Fig. 3 B). After genotyping (Supplementary Fig. 4 A, B), we confirmed the absence of centrioles and cilia in starved mESCs using immunofluorescence staining with antibodies against γ-tubulin and ARL13B (Fig. 3 C). All tested *Ift88* KO clones lacked primary cilia and in *Plk4* KO clones, both centrioles and cilia were lost. Given PLK4’s role in centriole duplication (Habedanck et al., 2005), we assessed whether its deletion affects cell proliferation. We, therefore, performed cell cycle analysis on all KOs for *Plk4* and *Ift88* by Propidium Iodide (PI) staining (Fig. 3 D). We observed no statistically significant variation between wild type and KO cell lines in their cell cycle profiles.

**Fig 3.**
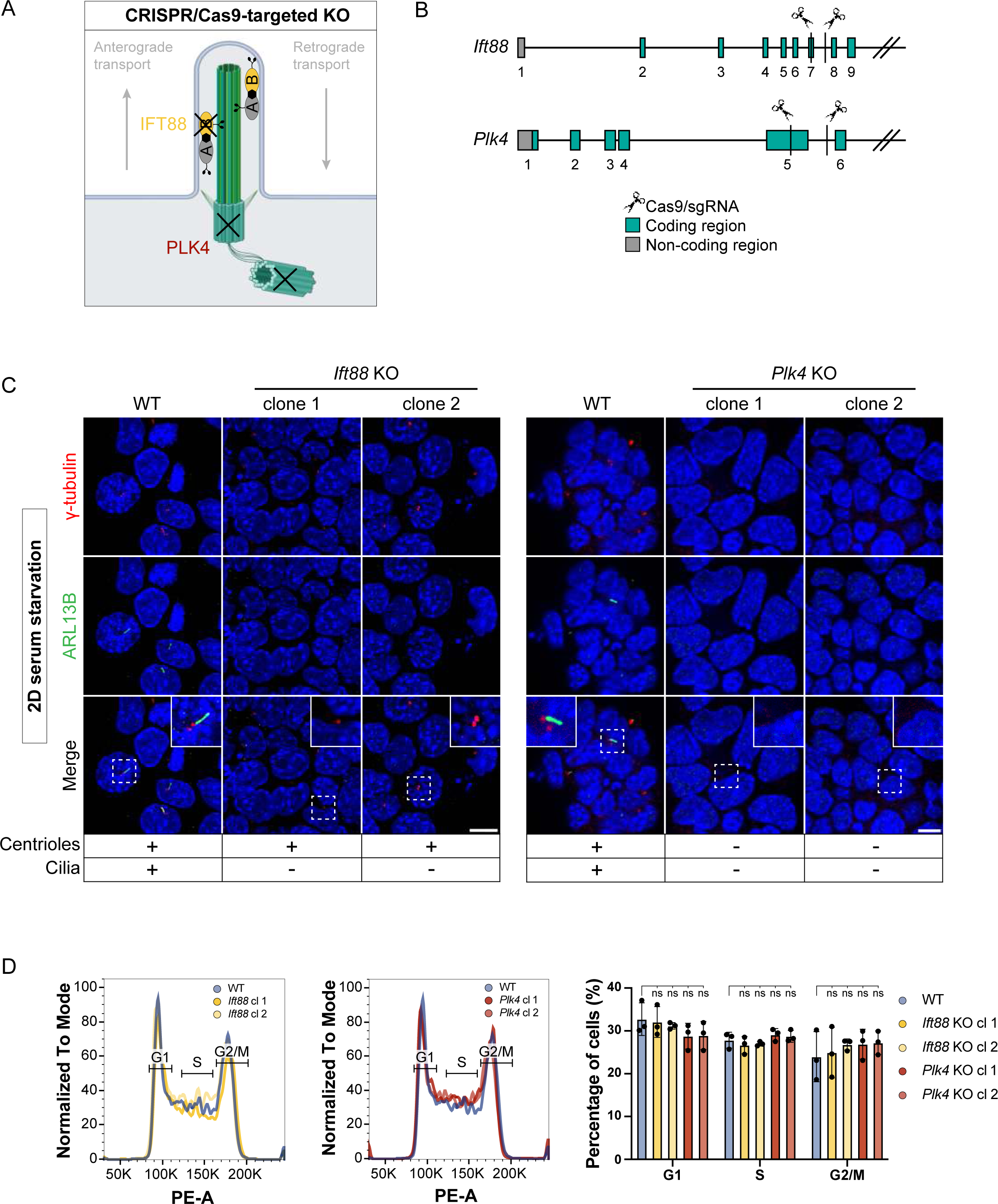
CRISPR KO of centriole/cilium components *Ift88* and *Plk4* depletes centrioles/cilia in 2D IF staining. (A) Schematic of a primary cilium indicating targets for CRISPR/Cas9 KOs. The cilium KO target IFT88 is a core component of the IFT-B train, while the centriole KO target PLK4 is a kinase essential for centriole duplication and hence indirectly also cilium formation. (B) Strategy for the generation of *Ift88* and *Plk4* KOs in mESCs. gRNAs are targeting exon 7 and the following intron in the *Ift88* KO, exon 5 and the following intron in the *Plk4* KO. Coding regions (green) and non-coding regions (gray) are indicated. (C) IF-validation of centriole and cilium KOs. Representative example of immunofluorescence staining of WT, *Ift88* and *Plk4* KO clones after induced ciliogenesis (48 h of starvation), labeled with antibodies against γ-tubulin (red), marker ARL13B (green) and DAPI staining (blue). Maximum intensity projection of central z-planes. (D) Cell cycle analysis by PI staining and flow cytometry of WT, *Ift88* and *Plk4* KO clones, n = 3 independent experiments (n.s.= non-significant, unpaired t test). Scale bar (C): 10 µm.

Taken together, we generated *Ift88* and *Plk4* KO cell lines using a CRISPR-targeted approach to remove centrioles and cilia from mESCs. These cell lines displayed a cell cycle profile comparable to wild type cells.

### 3D *in vitro* rosettes develop normally in the absence of centrioles and cilia

Polarization and lumenogenesis are key developmental processes during implantation associated with the re-organization of the epiblast (Kim et al., 2021). To study how the loss of centrioles or cilia affects these aspects, we generated 3D *in vitro* rosettes derived from either *Ift88* or *Plk4* KO clones. All *Ift88* and *Plk4* KO cell lines formed organized rosettes with a central lumen after 72 h of 3D rosette formation (Fig. 4 B). As expected, *Ift88* KO clones presented centrioles (γ-tubulin) but no cilia (ARL13B), while all *Plk4* KO clones showed neither centrioles nor cilia (Fig. 4 A). We quantified lumen and rosette size based on masking the central z-plane, using PODXL as a lumen marker and DAPI to assess rosette size. Measurements of the lumen area, rosette area and lumen-to-area ratio did not show significant differences between wild type and *Ift88* or *Plk4* KO rosettes (Fig 4 C).

**Fig 4.**
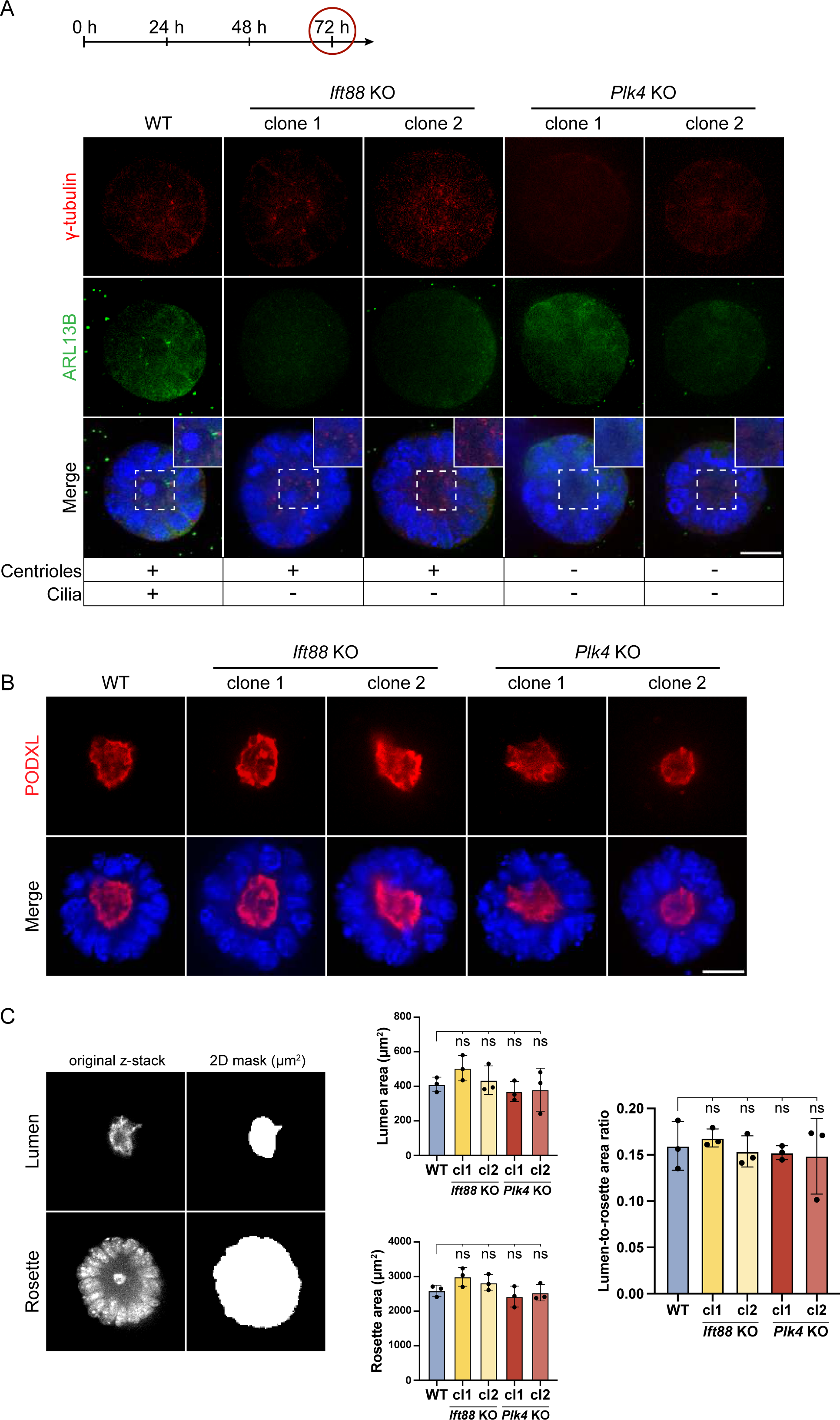
In *Ift88* and *Plk4* KO 3D rosettes, neither centrioles, nor cilia are required to form a lumen. (A) Validation of *Ift88* and *Plk4* KO clones in BME-embedded 3D *in vitro* rosettes growing without 2iLIF for 72 h. Representative immunofluorescence staining of rosettes labeled with antibodies against γ-tubulin (red), ARL13B (green) and DAPI staining (blue). The central z- plane is depicted. (B) Assessment of lumen formation of 3D BME-embedded *in vitro* rosettes after 72 h. Representative immunofluorescence staining labeled with antibodies against PODXL (red) and DAPI staining (blue). The central z-plane is depicted. (C) Quantification of lumen and rosette size based on the following masking model: 2D mask (area) of the central z-plane, using PODXL as a lumen marker and DAPI to assess rosette size. Column charts show lumen area, rosette area, and lumen-to-rosette area ratio of the central lumen and rosette z-planes. n = 3 independent experiments with 30 rosettes per condition/experiment (n.s.= non-significant, unpaired t test). Scale bar (A, B): 20 µm.

These findings suggest that centrioles and cilia are dispensable for polarization and lumenogenesis during *in vitro* 3D differentiation of early mouse development.

### Gastruloids recapitulate ciliogenesis *in vitro*

*Plk4* KO mouse embryos lack centrioles and arrest development at E7.5, characterized by delayed cell division and apoptosis (Hudson et al., 2001). Embryos lacking cilia generally do not survive beyond mid-gestation, around E11, largely due to failure of essential developmental signaling pathways dependent on ciliary function (Cortellino et al., 2009). However, loss of PLK4 or cilia did not affect early embryonic cell state transitions and morphogenesis *in vitro* (Figure 4 A, B). We therefore sought to investigate the role of centrioles and cilia in later stages of post-implantation development. In recent years, model systems have been developed for *in vitro* analysis of mammalian developmental progressions. However, to the best of our knowledge, these have not been applied to study ciliogenesis. Here, we implemented the mouse gastruloid assay, an *in vitro* model system that recapitulates early embryonic cell fate decisions until about 8.5 days after fertilization (Beccari et al., 2018), to study cilium and centrosome biology during development. In brief, 200 mESCs per well were aggregated for 48 h in differentiation media, followed by Wnt activation using a Chiron pulse for 24 h, leading to symmetry break, anterior-posterior polarization and germ layer differentiation (Fig 5 A, Supplementary Fig. 1 A, B). The protocol enables the cultivation of gastruloids for up to 120 h without shaking, corresponding up to E8.5 *in vivo*. We validated our system by verifying germ layer differentiation through spatiotemporal expression patterns of selected markers at different time points, ranging from 72 h to 120 h (Supplementary Fig. 1 A, B). The mesodermal marker Brachyury (Van Den Brink et al., 2014) was homogeneously expressed within gastruloids up to 72 h, consistent with global Wnt activation during the Chiron pulse. By 96 h, Brachyury prominently localized to the posterior end of the gastruloids, indicating symmetry break and the establishment of anterior-posterior polarity. Additionally, gastruloids were stained for the endoderm marker SOX17 (Kanai-Azuma et al., 2002) and SOX2, a marker expressed in neuro-mesodermal progenitors (NMPs), neural progenitors, and pluripotent cells (Avilion et al., 2003; Bergsland et al., 2011; Koch et al., 2017). We observed low and unpolarized SOX2 expression at 72 h. SOX2 signal gradually increased and accumulated at the posterior end of gastruloids from 96 h on. SOX17-positive cells were detected at the posterior pole of gastruloids, visible as a one-layered cell cluster surrounding a cavity. These findings are consistent with previous studies and show that in our hands gastruloid model effectively recapitulates symmetry break, anterior-posterior polarization and germ layer differentiation in the expected timeframe.

**Fig 5.**
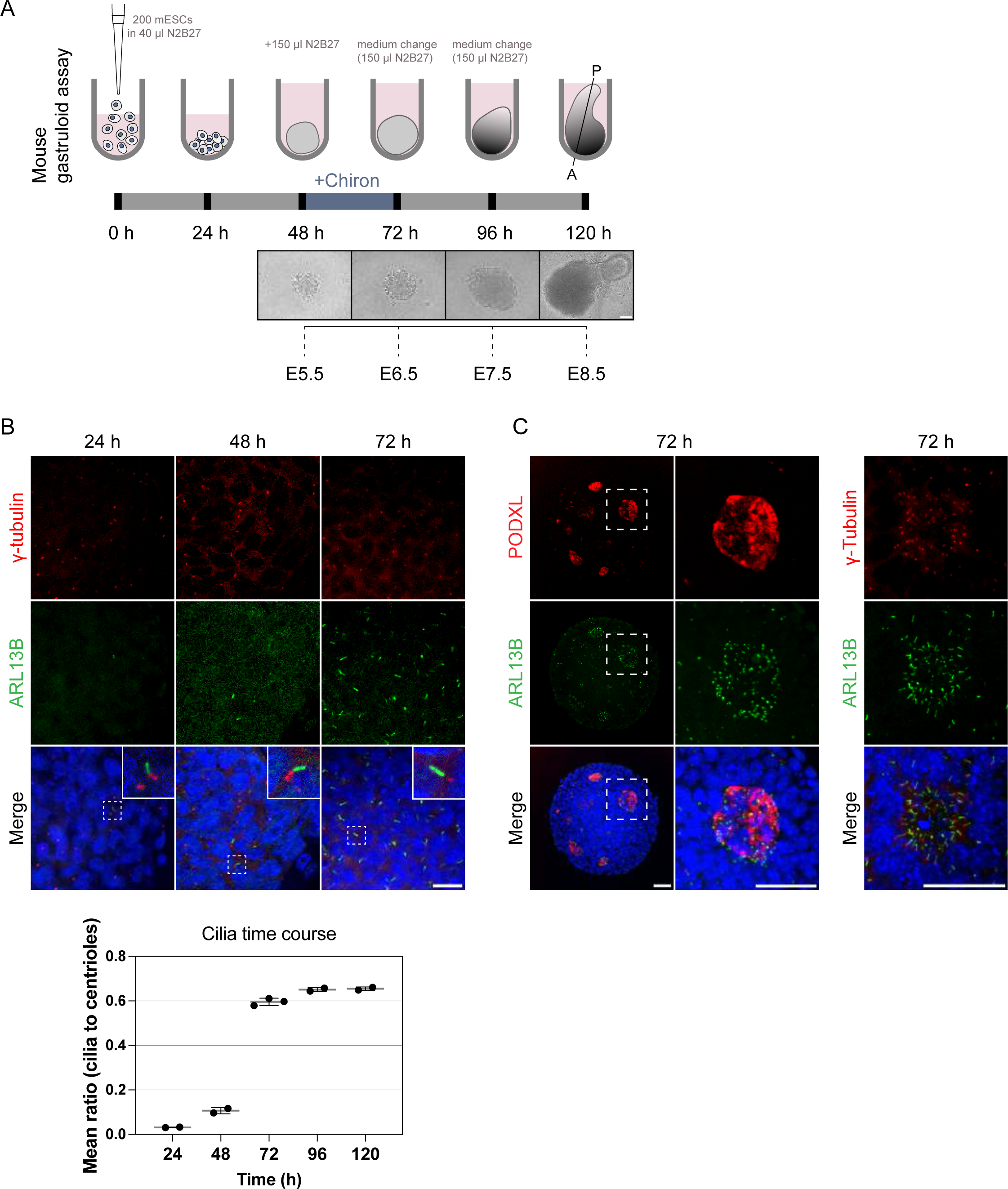
Gastruloids recapitulate early and late ciliogenesis *in vitro*. (A) Mouse gastruloid protocol up to 120 h of cultivation with representative brightfield images of different time points, compared to corresponding stages of *in vivo* mouse embryonic development. mESCs are cultivated in u-bottom, ultra-low attachment plates for 48 h in N2B27, followed by a Chiron pulse for 24 h, leading to symmetry break, anterior-posterior polarization and generation of all three germ layers, based on Beccari et al., 2018. (B) Representative example of immunofluorescence staining of gastruloids at 24, 48 and 72 h, labeled with antibodies against γ-tubulin (red), ARL13B (green) and DAPI staining (blue). Quantification of the mean ratio (cilia to centrioles) over a time course of 24 to 120 h, showing the increase of ciliogenesis during gastruloid development. Dots represent the mean ratio of individual experimental replicates per timepoint, with standard deviations indicated. (C) Lumenogenesis and ciliogenesis in gastruloids. Representative example of immunofluorescence staining of gastruloids at 72 h, labeled with antibodies against PODXL (left panel, red) or γ-tubulin (right panel, red), ARL13B (green) and DAPI staining (blue). Maximum intensity projection of central z-planes. Scale bar (A): 100 µm. Scale bar (B): 20 µm. Scale bar (C): 40 µm.

We next tested the suitability of gastruloids to study cilium formation during mammalian *in vitro* gastrulation. Gastruloids were fixed at 24, 48 and 72 h after seeding and immunostained with antibodies against ciliary markers (Fig. 5 B). Quantification of the mean cilia-to-centrioles ratio revealed sparse ciliation at 48 h, indicating that the majority of centrioles do not form cilia. A marked increase in ciliation was observed following the Chiron pulse at 72 h. Ciliated epithelial cells characteristically face a lumen (van der Vaart et al., 2021); therefore, we investigated whether this also occurs in gastruloids. Gastruloids cultured for 72 h effectively recapitulated key aspects of lumen formation, indicated by the lumen marker PODXL (Fig. 5 C). In line with our previous 3D rosette data (Fig. 2 D), primary cilia were enriched at cavities and projected into these luminal compartments.

Collectively, our results show that the gastruloid system is an ideal and versatile model system to implement in-depth studies of centrosome and cilia function during murine development by mimicking key developmental events including germ layer differentiation and anterior-posterior polarization.

### Cilia but not centrioles are dispensable in gastruloids

We next assessed the potential of *Plk4* and *Ift88* KO cells to form polarized, elongated mouse gastruloids. All tested *Plk4* KO clones displayed a severe phenotype, with cell aggregates progressively disassembling from the time of seeding onward, prior to the Chiron pulse (Fig. 6 A). By 72 h, the gastruloids completely degenerated and failed to progress further in development, potentially due to apoptotic events which will need to be further determined in future studies. We next investigated the role of cilia in gastruloids up to 120 h. Cilium depletion in the *Ift88* KO was confirmed in gastruloids, fixed and immunostained with antibodies against ARL13B at 120 h (Fig. 6 B). All tested *Ift88* KO clones continued to grow elongated anterior-posterior differentiated gastruloids up to 120 h (Fig. 6 C). We quantified gastruloid roundness and length based on the morphology of brightfield images (Supplementary Fig. 2 A). Both *Ift88* KO clones displayed a difference in gastruloid shape at 96 h, indicated by increased roundness and a decrease in length (Supplementary Fig. 2 B). Comparison of Brachyury and SOX2 in fixed and immunostained gastruloids revealed a similar spatiotemporal distribution of both markers in wild type and all tested *Ift88* KO clones at 72 and 120 h, with a marginal difference in tissue patterning of the *Ift88* KO at 96 h (Fig. 6 D).

**Fig 6.**
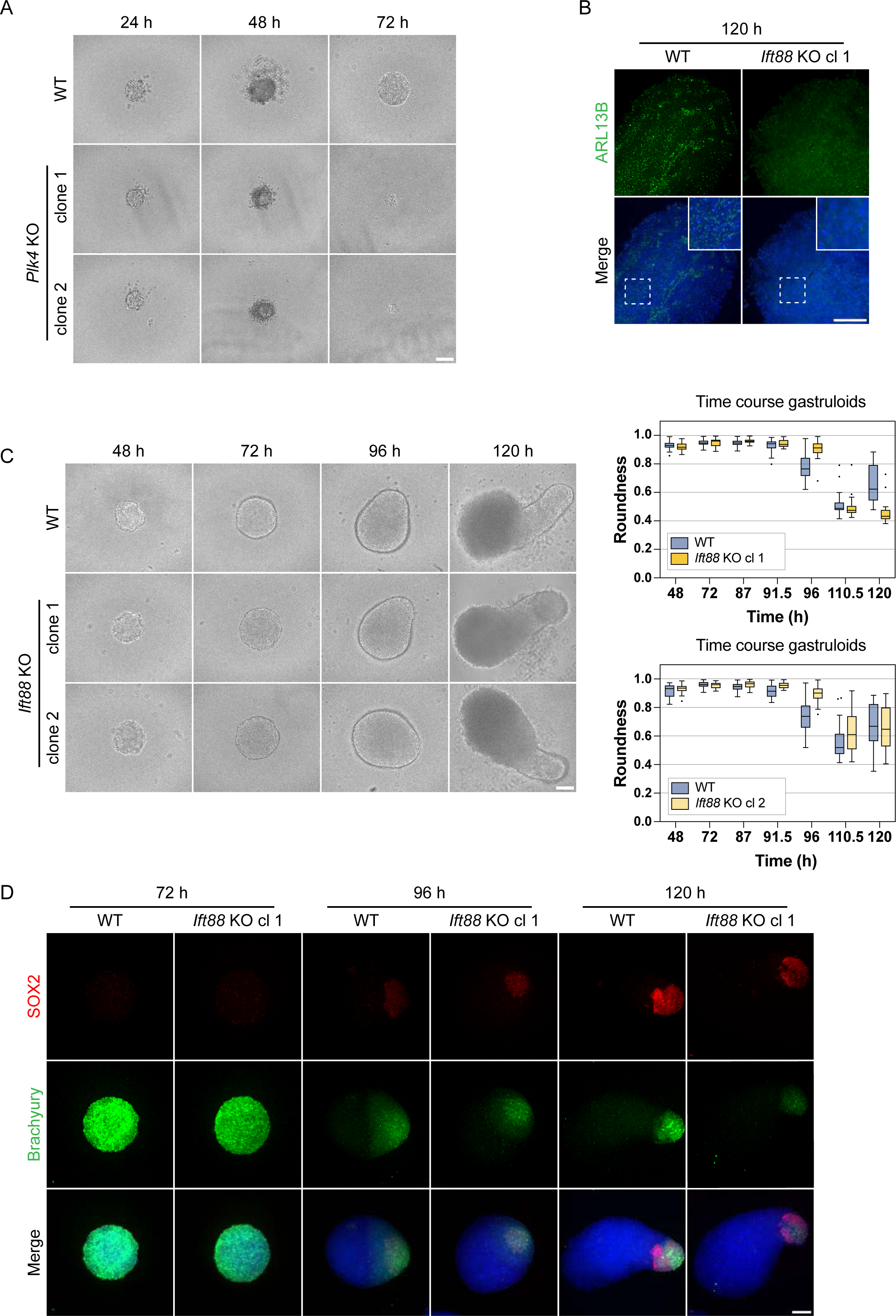
Gastruloids continue to elongate without cilia in the *Ift88* KO but disassemble in the centriole-depleted *Plk4* KO model. (A) Representative brightfield images of WT and *Plk4* KO gastruloids 24, 48 and 72 h after seeding. (B) Immunofluorescence staining of WT and *Ift88* KO gastruloids at 120 h, labeled with antibodies against ARL13B (green) and DAPI staining (blue). The maximum intensity projection is depicted. (C) Representative brightfield images of WT and *Ift88* KO gastruloids 48 to 120 h after seeding. Box plots indicating the roundness of WT and *Ift88* KO gastruloids. The box represents the interquartile range (IQR), with the median indicated by a horizontal line. Whiskers extend to 1.5 × IQR, and data points beyond this range are considered outliers (shown as dots). n=3 independent experiments with one representative experiment depicted. (D) Immunofluorescence staining of WT and *Ift88* KO gastruloids at 72, 96 and 120 h, labeled with antibodies against SOX2 (red), Brachyury (green) and DAPI staining (blue). The maximum intensity projection is depicted. Scale bar (A, B, C, D): 100 µm.

To conclude, centrioles are indispensable for gastruloid assembly, with loss of centrioles in *Plk4* KO clones resulting in gradual disassembly. In contrast, cilium-depleted *Ift88* KO clones continue to form gastruloids similar to the wild type and express mesoderm and neural progenitors, indicating proper anterior-posterior polarization, albeit with minor differences in gastruloid shape at 96 h.

### 3D *in vitro* gastruloids, generated from *Cep83^-/-^* and *Cep83Δexon4* cells, continue to develop successfully

In addition to our studies with *Plk4* and *Ift88* KO cell lines in different *in vitro* differentiation models, we dissected the function of the distal appendage protein CEP83 in early mouse development, using gastruloids as a model system (Fig. 7A). CEP83 is involved in anchoring the mother centriole to the cell membrane, a critical initiating step in mammalian ciliogenesis (Chong et al., 2020; Lo et al., 2019; Tanos et al., 2013). In contrast to *Ift88* depletion, which primarily impairs axoneme elongation, *Cep83* KO prevents the docking of the mother centriole to the plasma membrane, thereby completely abrogating ciliogenesis. This disruption also leads to the failure of IFT component recruitment and, consequently, loss of ciliary signaling (Joo et al., 2013). We, therefore, investigated the role of CEP83 in early mouse embryonic development using two different CRISPR/Cas9-KO strategies (Fig. 7 B). In KO strategy 1, we designed two gRNAs targeting exon 3 and 4, and in KO strategy 2, we targeted exon 4 along with the following intron. (Fig. 7 B). After genotyping via PCR (Supplementary Fig. 4 C, D), we designed primers to extract the *Cep83* cDNA both from the wild type and the different KO clones. The wild type showed the expected full-length *Cep83* construct while strategy 1 resulted in a frame shift generating multiple stop codons. However, the cDNA extracted from clones generated through KO strategy 2 consistently showed a version of *Cep83* where exon 4 was simply skipped and not included in the cDNA (Supplementary Fig. 5 C). We, therefore, labelled these clones *Cep83Δexon4.* We hypothesized that loss of this exon could be sufficient to lead to loss of cilia. Therefore, we validated the functional depletion of cilia in the mESC cell lines via immunofluorescence staining with antibodies against γ-tubulin and ARL13B. The cells were starved for 48 h to increase ciliogenesis for imaging experiments. All tested *Cep83^-/-^* and *Cep83Δexon4* clones failed to form cilia, as indicated by the absence of ARL13B expression (Fig. 7 C, D). We next evaluated the contribution of CEP83-depleted cell lines to generate polarized, elongated mouse gastruloids. All tested *Cep83^-/-^* and *Cep83Δexon4* clones continued to grow anterior-posterior elongated gastruloids up to 120 h (Fig. 7 E, F). Furthermore, all *Cep83^-/-^* and *Cep83Δexon4* clones displayed a difference in gastruloid shape before 120 h comparable to *Ift88* KO clones, most prominently at 96 h, indicated by increased roundness (Fig. 7 E, F) and a decrease in length (Supplementary Fig. 3 A, B). These data suggest that loss of cilia via *Ift88* or *Cep83* KO does not impair gastruloid formation, but delays gastruloid growth and elongation at around 96 h. To investigate whether this is also reflected in the spatiotemporal distribution of germ layer markers, we fixed gastruloids 72, 96 and 120 h after seeding and stained them with antibodies against Brachyury and SOX2. *Cep83^-/-^* and *Cep83Δexon4* gastruloids showed a similar spatiotemporal distribution of both markers compared to wild type at 72 h and 120 h, with less distinct foci at the posterior end at 96 h, similar to the *Ift88* KO (Supplementary Fig. 3 C, D). They expressed mesoderm and neural progenitors at the posterior end, indicating anterior-posterior polarization.

**Fig 7.**
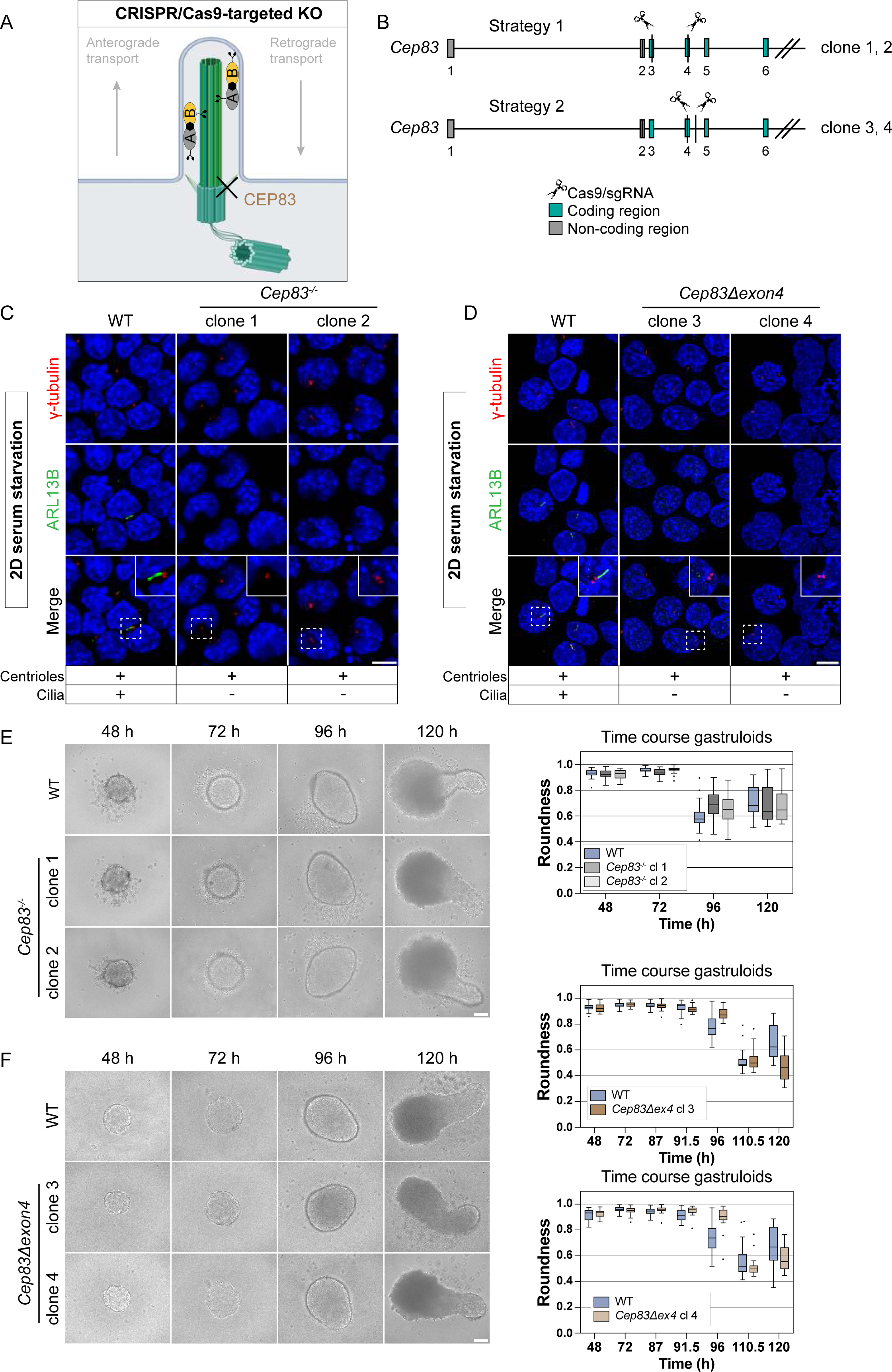
*Cep83* is not essential for gastruloid formation. (A) Scheme of a primary cilium indicating the CRISPR/Cas9 KO target *Cep83*. CEP83 is a distal appendage protein (DAP), required for anchoring the mother centriole to the plasma membrane and hence for ciliogenesis. (B) Strategy for the *Cep83* KO generation in mESCs. gRNA pairs are targeting exon 3 and 4, and exon 4 and the following intron. Coding regions (green) and non-coding regions (gray) are indicated. (C, D) IF-validation of cilium depletion in *Cep83^-/-^* and *Cep83Δexon4* clones. Representative example of immunofluorescence staining of WT, Cep83^-/-^ and *Cep83Δexon4* clones after induced ciliogenesis (48 h of starvation), labeled with antibodies against γ-tubulin (red), ARL13B (green) and DAPI staining (blue). Maximum intensity projection of central z-planes. E) Representative brightfield images of WT and *Cep83^-/-^*gastruloids 48 to 120 h after seeding. Box plot showing the roundness of WT and *Cep83^-/-^* gastruloids. The box represents the interquartile range (IQR), with the median indicated by a horizontal line. Whiskers extend to 1.5 × IQR, and data points beyond this range are considered outliers (shown as dots). n=3 independent experiments with one representative experiment depicted. (F) Representative brightfield images of WT and *Cep83Δexon4* gastruloids 48 to 120 h after seeding. Box plot showing the roundness of WT and *Cep83Δexon4* gastruloids. The box represents the interquartile range (IQR), with the median indicated by a horizontal line. Whiskers extend to 1.5 × IQR, and data points beyond this range are considered outliers (shown as dots). n=3 independent experiments with one representative experiment depicted. Scale bar (C, D): 10 µm. Scale bar (E, F): 100 µm.

In summary, our findings demonstrate that the depletion of cilia neither through *Ift88* KO nor through the loss of *Cep83* disrupts gastruloid differentiation.

### Naive *Cep83Δexon4* mESCs, initially devoid of cilia, successfully regenerate cilia in rosette and gastruloid differentiation

The comparison of 2D cultures to more sophisticated 3D systems frequently highlights discrepancies between these culture systems with relevance of context-dependent tissue architecture. Fundamental distinctions include extracellular matrix interactions, nutrient availability, and cellular polarization, which are essential features of the 3D microenvironment lacking in 2D cultures (Duval et al., 2017; Kapałczyńska et al., 2018). Since *Cep83Δexon4* clones still express a truncated form of CEP83, we conducted a reassessment of cilium depletion of *Cep83^-/-^* and *Cep83Δexon4* clones in gastruloids. As expected, *Cep83^-/-^* gastruloids did not exhibit cilia after 120 h (Fig. 8 A). However, *Cep83Δexon4* gastruloids expressed rod-shaped ciliary structures indistinguishable from wild type, with a γ-tubulin and ARL13B localization pattern and ciliary length (∼ 2 µm) comparable to the wild type (Fig. 8 B). Time course experiments indicated the gradual initiation of ciliogenesis in *Cep83Δexon4* gastruloids, beginning at 72 h - a time point at which ciliogenesis increased in the wild type - albeit at initially lower levels compared to the wild type (Fig. 8 C). However, by 120 h, cilia levels in the mutant reached those observed in the wild type. This finding was unexpected, as cilia were consistently absent in undifferentiated *Cep83Δexon4* mESCs in 2D (Fig. 7 C, D). We investigated whether *Cep83Δexon4* acquires a functional role during 3D gastruloid formation and whether this effect is dependent on its expression levels. To determine whether the lack of cilia in *Cep83Δexon4* mESCs and the delayed initiation of ciliogenesis in *Cep83Δexon4* gastruloids can be rescued by *Cep83Δexon4* overexpression, we designed *Flag-Cep83/Cep83Δexon4* overexpression mESC lines. Immunostaining showed that overexpression of full-length CEP83 in the *Cep83Δexon4* cell line successfully rescued cilium formation (Supplementary Fig. 5 A). However, overexpression of CEP83Δexon4 failed to restore ciliogenesis, suggesting that CEP83Δexon4 lacks functional activity in 2D mESCs, even upon overexpression. We next investigated whether CEP83Δexon4 overexpression can rescue the delayed onset of ciliogenesis in gastruloids (Supplementary Fig. 5 B). Quantification of cilia in gastruloids fixed at 72 h indicated the most prominent difference in ciliation between wild type and *Cep83Δexon4* cell lines. Overexpression of truncated CEP83Δexon4 in the *Cep83Δexon4* cell line partially rescued cilium formation but failed to restore cilium levels similar to the wild type. In contrast, overexpression of full-length CEP83 in CEP83Δexon4 gastruloids fully restored cilium levels comparable to wild type. These findings suggest that CEP83Δexon4 overexpression can partially rescue ciliation in mouse gastruloids at 72 h; however, it is not reaching wild type levels.

**Fig 8.**
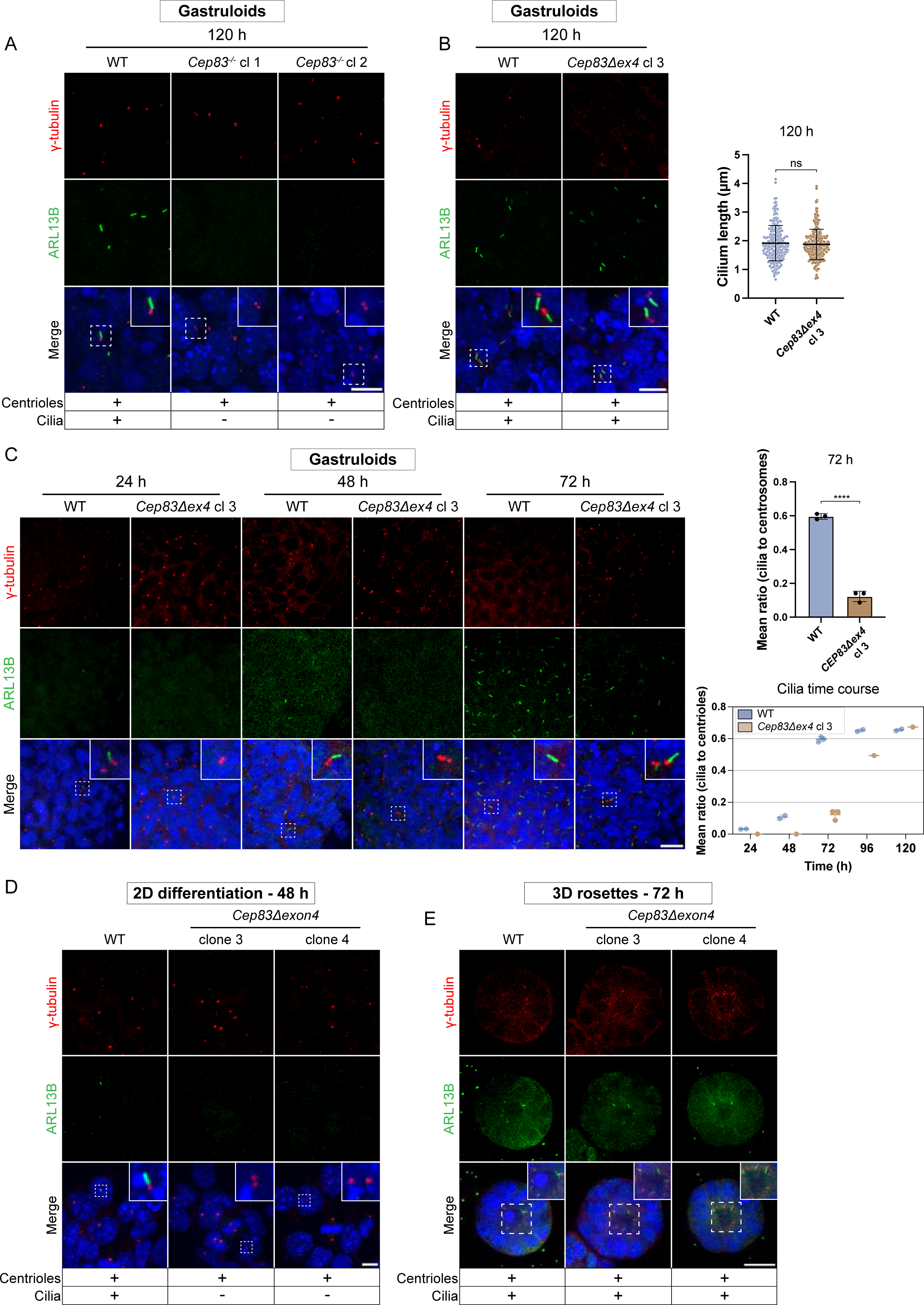
Cilium-depleted, naive *Cep83Δexon4* mESCs regain cilia in 3D rosettes and gastruloids. (A) Immunofluorescence staining of WT and *Cep83^-/-^* gastruloids at 120 h, labeled with antibodies against γ-tubulin (red), ARL13B (green) and DAPI staining (blue). The maximum intensity projection of the 5 central z-planes is depicted. (B) Immunofluorescence staining of WT and *Cep83Δexon4* gastruloids at 120 h, labeled with antibodies against γ-tubulin (red), ARL13B (green) and DAPI staining (blue). The central z-plane is depicted. Cilium length (µm) was quantified in at least 229 cilia per condition (mean + SD, n.s.= non-significant, unpaired t test). (C) Representative immunofluorescence staining of WT and *Cep83Δexon4* gastruloids at 24, 48 and 72 h, labeled with antibodies against γ-tubulin (red), ARL13B (green) and DAPI staining (blue). Quantification of the mean ratio of cilia to centrioles in WT and *Cep83Δexon4* after 72 h (****p < 0.0001, unpaired t test). Quantification time course of the mean ratio of cilia to centrioles from 24 to 120 h. Dots represent the mean ratio of individual experimental replicates per timepoint, with standard deviations indicated. (D) Immunofluorescence staining of formative EpiLCs after 48 h of differentiation labeled with antibodies against γ-tubulin (red), ARL13B (green) and DAPI staining (blue). Maximum intensity projection is depicted. (E) Validation of *Cep83Δexon4* clones in BME-embedded 3D *in vitro* rosettes growing without 2iLIF for 72 h. Representative immunofluorescence staining labeled with antibodies against γ-tubulin (red), ARL13B (green) and DAPI staining (blue). The central z-plane is depicted. Scale bar (A, B, D): 10 µm. Scale bar (C, E): 20 µm.

To understand why exon 4 of *Cep83* becomes dispensable for ciliogenesis in gastruloids, we investigated the reliance of cilium restoration on exit of naive pluripotency by differentiating mESCs into EpiLCs in 2D culture for 48 h. Cilia were absent in these differentiated *Cep83Δexon4* clones, suggesting that this phenomenon is not driven by the transition out of naive pluripotency, but rather by an alternative mechanism (Fig. 8 D). We hypothesized that cell polarization, establishment of the extracellular matrix and morphogens, present in a 3D environment, enable ciliogenesis independent of exon 4 of *Cep83*. To test this, we differentiated *Cep83Δexon4* cells into 3D *in vitro* rosettes, which polarize and form lumen during exit of naive pluripotency but without differentiating further into all germ layers. 3D rosettes, derived from *Cep83Δexon4* clones, expressed cilia after 72 h (Fig. 8 E). These data suggest that the truncated isoform of CEP83 is non-functional under 2D conditions. In contrast, *Cep83Δexon4* cells grown in 3D rosettes or gastruloids form cilia, indicating that the CEP83Δexon4 variant is able to bypass potential limitations encountered in 2D. However, the mechanism enabling CEP83Δexon4 activity in differentiation remains unclear and requires further investigation in the future.

In summary, we utilize different 3D model systems that serve as novel tools to study ciliogenesis in early embryonic development, from implantation to gastrulation, recapitulating not only key developmental events but also context-specific cilium assembly. Our study highlights the importance of incorporating complex 3D environment setups in developmental studies and their advantages over less advanced 2D systems.

## Discussion

Primary cilia and centrioles play essential roles in mammalian embryonic development, yet their precise contributions to embryonic signaling and morphology in the early embryo before the formation of the node remain incompletely understood. In mice, centrioles are initially absent in the fertilized embryo and first assemble *de novo* at the blastocyst stage at E3.5 (Courtois et al., 2012; Gueth-Hallonet et al., 1993; Howe & FitzHarris, 2013). Yet, only a short time later they are clearly functionally important, with their loss resulting in embryonic arrest by E7.5 (Hudson et al., 2001). Meanwhile, primary cilia first arise on epiblast cells during cavitation at E5.5-E6 (F. K. Bangs et al., 2015), and ciliary mutants exhibit mid-gestation arrest (E11) due to defects in Hh-dependent neural and limb patterning (Cortellino et al., 2009; Huangfu et al., 2003). It is essential to understand the processes occurring between the initiation of centriole and cilium formation and the onset of embryonic lethality to elucidate their developmental roles. In this study, we used different 3D *in vitro* model systems to study centrioles and cilia in early mouse embryonic development and their contribution to polarization, germ layer differentiation, and symmetry breaking.

Recent advances in cilium research highlight the close interplay between cell differentiation and ciliogenesis (Yanardag & Pugacheva, 2021), with primary cilia acting as critical signal transducers that regulate cell differentiation in various tissues (Arrighi et al., 2017; Coschiera et al., 2024; Shim et al., 2023). We investigated whether cell differentiation *per se* can promote cilium formation. The widely used transition from mESCs to EpiLCs reflects many transcriptional changes observed in the early mouse embryo; however, this transition did not enhance ciliogenesis, suggesting that increased primary cilium formation is not an inherent feature of exit from naive pluripotency. In the 3D rosette model system, where the differentiating mESCs are embedded into extracellular matrix, cells similarly exit from naive pluripotency, while also undergoing morphological changes including polarization and lumenogenesis (Bedzhov & Zernicka-Goetz, 2014; Shahbazi et al., 2017). Under these conditions, the differentiating mESCs exhibit increased ciliation, demonstrating further that ciliation is not coupled to differentiation *per se* but also to the 3D microenvironment.

We generated centriole (*Plk4*) and cilium (*Ift88*) KO cell lines, to study the function of centrioles and cilia in the 3D *in vitro* rosette model system. Loss of *Plk4* and *Ift88* did not disrupt polarization and lumenogenesis, with the establishment of well-organized rosettes of similar size after 72 h of 3D rosette culture. Our results demonstrate that centrioles and cilia are not required for 3D *in vitro* differentiation and lumenogenesis during early mouse development, consistent with the reported embryonic arrest of *Plk4^-/-^* mice only after gastrulation (Hudson et al., 2001; Swallow et al., 2005).

While 3D rosettes are a great model system to study cell state changes during the implantation stage, cells will not undergo further developmental changes normally observed during gastrulation. Therefore, we employed mouse gastruloids, a powerful model system recapitulating germ layer specification and *in vivo* development up to E8.5 (Beccari et al., 2018; Hashmi et al., 2022; Stelloo et al., 2024). *Plk4* KO mouse gastruloids disassembled from the time of seeding onward, prior to the Chiron pulse, and failed to progress further in development. Thus far, it remains unclear whether cells simply dissociate or are actively eliminated, potentially mediated by p53 activation. Centrosomal activity is essential for cell cycle progression (Hinchcliffe et al., 2001; Khodjakov & Rieder, 2001). Several studies have shown that loss of centrioles results in cell cycle arrest and p53-dependent apoptosis (Bazzi & Anderson, 2014; Grzonka & Bazzi, 2024; Lambrus et al., 2015; Wong et al., 2015). One proposed mechanism causing this arrest is the increased duration of mitosis in centriole-depleted mutant mouse embryos (Bazzi & Anderson, 2014). The duration of mitosis is monitored during cell cycle progression; extended mitosis, including cumulatively over multiple cell cycles, has been associated with G1 arrest, an ability that is often lost in p53-mutant aberrant cells (Meitinger et al., 2024). Interestingly, in *Drosophila melanogaster*, a delay in mitotic spindle assembly following loss of centrioles has also been observed, with the key difference that these embryos develop to the adult stage without reported cell death and only die postnatally due to the absence of cilia (Basto et al., 2006). This suggests that vertebrates possess more stringent cell cycle checkpoints to eliminate acentriolar cells. Although the observed disassembly in our *Plk4* KO gastruloids resembles embryonic arrest of *in vivo* data at E7.5 (Hudson et al., 2001), the onset of the phenotype occurs earlier than expected, with gastruloids gradually disassembling from 48 h on, roughly corresponding to E5.5-E6.5. The discrepancy between the timing of the phenotype observed *in vitro* and *in vivo* could be attributed to the gastruloid model system itself, which is derived from mESCs that possess fully functional and matured centrioles, whereas these are typically absent until the blastocyst stage *in vivo* (Courtois et al., 2012; Gueth-Hallonet et al., 1993; Howe & FitzHarris, 2013). This difference may account for the accelerated onset of the phenotype in our model. In addition to its role in centriole duplication, PLK4 has been reported to have a centriole-independent function in early mouse embryos, where it initiates acentriolar spindle assembly in mammalian oocytes (Bury et al., 2017; Coelho et al., 2013). However, our data show that the loss of Plk4 has no effect on cell viability in mESCs.

Whereas centriole depletion significantly impaired gastruloid formation, cilia loss in *Ift88* KO mESCs did not perturb the formation of elongated anterior-posterior differentiated gastruloids, exhibiting a spatiotemporal expression of mesoderm (Brachyury) and neuronal progenitor cells (SOX2) similar to the wild type. It has been previously demonstrated that cilium KO mouse embryos undergo arrest at mid-gestation around E11 (Cortellino et al., 2009; Huangfu et al., 2003). The absence of a severe phenotype in *Ift88* KO gastruloids may be attributed to the fact that they can only be cultured up to 120 h without shaking, thereby recapitulating developmental stages only from E5.5 to E8.5 as previously described (Arias et al., 2022; Beccari et al., 2018; Stelloo et al., 2024).

Interestingly, although *Ift88* KO gastruloids were able to elongate up to 120 h, they exhibited a difference in morphology at 96 h, appearing rounder and less elongated compared to controls. This effect was consistently observed across all tested clones that lacked cilia while it was not observed in other, unrelated KO clones (data not shown). In wild type gastruloids, only a small fraction of cells is ciliated up to 48 h. The number increases sixfold after 72 h, corresponding to the time frame cilia first appear in the mouse embryo at E5.5–E6.5 (F. K. Bangs et al., 2015). After 72 h, ciliogenesis reached a plateau and remained stable throughout the culture period until 120 h. The timing of the potential delay in gastruloid elongation coincides with the peak of ciliogenesis in wild type gastruloids and may be attributed to the absence of ciliary signaling. Hh signaling, for example, has been shown to be impaired in cells with disrupted IFT, leading to abnormal limb and neural tube patterning (Haycraft et al., 2005; Huangfu et al., 2003; Huangfu & Anderson, 2005; Liu et al., 2005). Additionally, Wnt signaling is involved in different aspects of development, including establishment of an anterior-posterior axis, cell fate decisions, proliferation and cell death (Vuong & Mlodzik, 2023). Whether the loss of ciliary signaling pathways in *Ift88* KO gastruloids underlies the observed morphological changes and why this phenotype does not persist up to 120 h remains to be determined. Our gastruloid cultivation is limited to 120 h, after which, and occasionally even earlier, disassembly begins. The loss of phenotype might therefore reflect a general loss of gastruloid integrity rather than the existence of compensatory mechanisms. Distinguishing between these possibilities would require conditions allowing gastruloids to be cultured beyond 120 h using alternative long-term cultivation protocols.

In addition to our studies with *Plk4* and *Ift88* KO cell lines in different *in vitro* model systems, we investigated the role of the centriolar distal appendage protein CEP83 in early mouse development. Distal appendages are involved in anchoring the mother centriole to the cell membrane (Chong et al., 2020; Lo et al., 2019; Tanos et al., 2013), and *Cep83* KO completely inhibits ciliogenesis, preventing IFT recruitment and resulting in a loss of ciliary signaling (Joo et al., 2013). As anticipated, our data demonstrate that *Cep83^-/-^* cell lines are capable of generating polarized, elongated mouse gastruloids, similar to what was observed in *Ift88* KO clones. Limited evidence indicates that ciliogenesis might still take place in CEP83 mutants, although it appears to be less efficient and is frequently associated with structural defects (Mansour et al., 2022; Shao et al., 2020). In our study, *Cep83^-/-^* cells lacked cilia, whereas the truncated isoform *CEP83Δexon4*, unexpectedly recovered ciliogenesis during 3D differentiation in gastruloids. Ciliation occurred gradually in gastruloids and reached wild type levels at 120 h. This suggests that exon 4 of CEP83 is important for ciliogenesis in non-polarized cells, such as during 2D cultivation of mESCs, but is only partially required during 3D differentiation. While overexpression of the *Cep83Δexon4* variant did not rescue ciliogenesis in 2D, overexpression of the isoform in gastruloids partially rescued ciliation, revealing a context-dependent function of exon 4. The main difference between 2D and 3D cultivation is the polarization of the cells. Therefore, it is tempting to speculate that in the absence of polarization, truncated CEP83 cannot anchor the basal body to the membrane. Regardless of the precise mechanism, this discrepancy highlights the importance of studying cellular processes in their proper 3D context, with conventional 2D cultures potentially yielding misleading results.

In conclusion, our study presents the first *in vitro* study of centriole and cilium formation in early mouse development, employing different 3D models to recapitulate key events of implantation, tissue patterning and anterior-posterior elongation. Although these models do not represent true embryos, they provide valuable platforms for investigating centriole and cilium formation and function within a highly controlled and closely monitored experimental framework.

## Methods

### Cell culture maintenance

Male R1 mESCs (Nagy et al., 1993) were routinely cultivated at 37°C, 5% CO2 in “2iLIF medium”, composed of N2B27 base medium - HyClone™ DMEM/F12 1:1 mix (Cytiva, SH30271.FS) with 2.5 mM L-Glutamine and without HEPES, 4 g/L AlbuMAXTM II (Gibco™, 11021029), 1x serum-free B-27™ Supplement (Gibco™, 17504044), 1x N2 components (homemade, Sigma-Aldrich, R&D Systems), 1x MEM NEAA (Gibco™, 11140035), 1x Penicillin-Streptomycin (Gibco™, 15070063), 1x Sodium Pyruvate (Gibco™, 11360039), 0.055 mM 2-Mercaptoethanol (Gibco™, 21985023), supplemented with the MEK inhibitor Mirdametinib (PD0325901, 0.8 µM, MedChemExpress), the GSK3β inhibitor Laduviglusib (CHIR99021, 3.3 µM, MedChemExpress) and 10 ng/mL human LIF (provided by the VBCF Protein Technologies Facility, https://www.viennabiocenter.org/facilities/) on CytoOne® Multi-Well Plates. Prior to usage, tissue culture dishes were pre-coated with 7.5 µg/ml Poly-L-ornithine hydrobromide (Sigma-Aldrich, P4638) for 1 h at 37°C, followed by 5 µg/ml Laminin (Sigma-Aldrich, L2020) for 1 h at 37°C. The cells were passaged every two to three days in an appropriate ratio using 1x Trypsin-EDTA solution (Sigma-Aldrich, T3924) for 3 min at 37°C to detach the cells and 10% Fetal Bovine Serum (Sigma-Aldrich, F7524) to stop the reaction. Cells were regularly tested for mycoplasma contamination using the MycoAlertTM PLUS Mycoplasma Detection Kit (Lonza, LT07-705).

### mESC to EpiLC differentiation (2D)

For differentiation of naive mESCs to formative EpiLCs, 10 000 cells/well were seeded in N2B27 without 2iLIF on µ-Slide 8 Well Glass Bottom Plates (ibidi, 80827) coated with fibronectin (YO Proteins, #663, 10µg/ml). Undifferentiated controls were seeded in parallel and cultured in 2iLIF. After 48 h of growth, cells were fixed and stained for immunofluorescence microscopy according to the protocol below (Fixation and immunofluorescence staining of 2D cultured cells).

### Cell starvation (2D)

10 000 cells were seeded in 2iLIF on fibronectin-coated (10 µg/mL) µ-Slide 8 Well Glass Bottom Plates (ibidi, 80827). After 24 h, cells were washed once with 250 µl Dulbecco′s Phosphate Buffered Saline (PBS, Sigma-Aldrich, D8537) and the starvation was initiated by addition of 250 µl starvation medium (DMEM/F12, supplemented with 0.1% Fetal Bovine Serum (FBS), 0.8 µM MEK inhibitor PD0325901 and 3.3 µM GSK3β inhibitor CHIR99021). Cells were starved for 24 to 48 h, as indicated in the figure legends. The controls remained in 2iLIF for the respective time. Samples were fixed and stained for immunofluorescence microscopy according to the protocol below (Fixation and immunofluorescence staining of 2D cultured cells).

### Fixation and immunofluorescence staining of 2D cultured cells

2D cultured cells were 1x washed with PBS and fixed with 250 µl PFA (pre-warmed 37°C) for 15 min at room temperature. After 3x washing with PBST (PBS + 0.1 % TWEEN® 20), cells were permeabilized with 250 µL 0.1% Triton-X/PBS for 10 min at room temperature, washed 3x with PBST and incubated for 30 min in 250 µl blocking buffer (PBST + 5% BSA) at room temperature. Next, cells were stained with primary antibody (diluted in blocking buffer) overnight at 4°C. After 3x washing with PBST, cells were incubated with secondary antibody solution for 1 h at room temperature (negative control received only the secondary antibody). Finally, samples were incubated in DAPI solution (1 µg/ml, Sigma-Aldrich, D9542) for 10 min, washed 3x in PBS and stored at 4°C until image acquisition.

### 3D *in vitro* rosette assay

Previously described 3D rosette formation assay (Bedzhov & Zernicka-Goetz, 2014; Shahbazi et al., 2017) was adjusted and optimized for our R1 mESC line to stably recapitulate polarization, lumenogenesis and cilium formation during mouse implantation. mESCs were cultivated in 2iLIF at 37°C, 5% CO2 and split at least 1x prior to the rosette formation assay to adjust to our culture conditions. Cells were trypsinized, washed two times in PBS, pelleted (20 000 cells/pellet) and resuspended in 20 µl ice-cold Cultrex Reduced Growth Factor Basement Membrane Extract (BME), Type 2 (R&D Systems, 3533-005-02), which mimics the basement membrane that surrounds the epiblast during implantation (Bedzhov & Zernicka-Goetz, 2014). Each 20 µl cell suspension drop was carefully placed in the center of a µ-Slide 8 Well Glass Bottom Plate (ibidi, 80827) and incubated for 10 min at 37°C to enable the BME to solidify. 250 µl of pre-warmed N2B27 base medium or 2iLIF medium (depending on the assay) was added to each well and plates cultivated at 37°C in 5% CO2 for up to 72 h with daily medium changes.

### Fixation and immunostaining of rosettes

3D rosettes were washed with 1x PBS and fixed with 250 µl (pre-warmed 37°C) 4% paraformaldehyde (PFA) (methanol free, MP Biomedicals, 0219998380) for 30 min at room temperature. Cells were washed 3x with PBST (0.1% TWEEN in PBS) and permeabilized with 250 µL 0.3% Triton-X/PBS for 30 min at room temperature. After washing 3x with PBST, 250 µl blocking buffer was added to each well and incubated for 30 min at room temperature. Next, samples were stained with primary antibodies (diluted in blocking buffer) against Podocalyxin (R&D Systems, MAB1556) for lumen quantifications, ARL13B (Proteintech, 17711-1-AP) and γ-tubulin (Sigma-Aldrich, T6557) for KO validations, at 4°C overnight. After 3x washing with PBST, the secondary antibody Anti-Rat IgG H&L Alexa Fluor® 555 (abcam, ab150154) for lumen quantifications, Anti-Rabbit Alexa Fluor® 488 (abcam, ab150073) and donkey Anti-Mouse Alexa Fluor® 555 (abcam, ab150106) for KO validations, incubated for 1 h at room temperature, including the negative control, containing only the secondary antibody. Finally, samples were incubated in DAPI solution (1 µg/ml) for 15 min, washed 3x in PBS and stored at 4°C until image acquisition.

### Gastruloid culture

mESCs were cultivated up to 80% confluency in 2iLIF at 37°C, 5% CO2 and passaged 2x before starting the gastruloid assay to provide stable cell growth and ensure cell adaptation to medium conditions. The gastruloid differentiation protocol was previously described (Beccari et al., 2018). mESCs were detached with trypsin, washed 3x with PBS and resuspended in gastruloid N2B27 medium – 50% DMEM/F12 with GlutaMAX™ (Gibco™, 31331028), 50% Neurobasal™ medium (Gibco™, 21103049), 1x GlutaMAX™ (Gibco™, 3133102), 1x MEM NEAA (Gibco™, 11140035), 1x Penicillin-Streptomycin (Gibco™, 15070063), 1x Sodium Pyruvate (Gibco™, 11360039), 0.1 mM 2-Mercaptoethanol (Gibco™, 21985023), 1x N-2 Supplement (Gibco™ 17502048) and serum-free B-27™ Supplement (Gibco™, 17504044). 200 cells were seeded per well in 40 µl gastruloid N2B27 medium in Corning™ 96-Well Clear Ultra Low Attachment Microplates (Corning™, 7007) and incubated at 37°C, 5% CO2. 48 h after seeding, 150 µl of gastruloid N2B27 medium with 3 µM GSK3β inhibitor CHIR99021 was added to each well. After 24 h, the Chiron-pulse was stopped by replacing 150 µl of Chiron-medium with 150 µl of basic gastruloid N2B27 medium. The medium was changed every day and images acquired using the ZOE Fluorescent Cell Imager (Bio-Rad).

### Gastruloid fixation and immunostaining

Gastruloids were prepared for imaging as previously described (Beccari et al., 2018). Prior to the staining procedure, plates were coated in gastruloid blocking buffer PBS-FT (PBS, 10% FBS, 0.2% Triton) to avoid attachment of gastruloids. All pipette tips were cut and pre-coated in PBS-FT solution to ensure safe gastruloid transfer without damage or loss. Gastruloids differentiated for 48, 72 and 96 h in Corning™ 96-Well Clear Ultra Low Attachment Microplates (Corning™, 7007) were then transferred to a 6-well plate with 2 ml PBS. All replicate wells of the same condition were pooled in one well. To enable easier transfer, the plate was moved in a circular manner until all gastruloids accumulated in the center of each well. They were collected with a 1 ml pipette, kept in a vertical position to enable gastruloid sedimentation at the bottom of the tip. The tip was slightly pushed and the “hanging drop”, containing mainly gastruloids with minimal carry-over, was transferred to a new well containing 2 ml of 4% PFA. Samples were incubated for 2 h at 4°C. Gastruloids were washed 3x with PBS-FT, each washing step containing a transfer step into a new well and remained for 10 min in the last well. Samples were either stored at 4°C until further use or the staining procedure was continued. They were transferred to a 12-well plate and blocked in 2 ml PBS-FT for 1 h on an orbital shaker. In order to reduce antibody volumes, gastruloids were placed in a 48-well plate containing 120 µl primary antibody solution and incubated for 24 h on an orbital shaker at 4°C. Samples were washed 3x with PBS-FT, including a 20 min incubation time of the last washing step and stained in 120 µl secondary antibody solution and DAPI staining (2.5 µg/ml) while shaking overnight at 4°C. After washing gastruloids 3x in PBS-FT, they were mounted. Cleaning of microscope slides with ethanol ensured removal of dust particles. A drop of 20 µl of VECTASHIELD® Antifade Mounting Medium with DAPI (Vector Laboratories, VEC-H-1200) was placed to the center of a microscope slide and a small drop to the center of the cover slip to avoid air bubbles. Up to 10 gastruloids (depending on the time point and their size to avoid clustering) were placed within each drop on the microscope slide and carefully covered with a cover slip. After removal of excess liquid, the border of the cover glass was sealed with several layers of nail polish and all specimens stored at 4°C until imaging.

### Generation of KO cell lines

Cilium and centriole KO cell lines were generated with CRISPR/Cas9 (Cong et al., 2013). Forward and reverse DNA oligonucleotides were designed in Benchling, containing the gRNA-Sequence to the target gene as well as the overhangs required for cloning and were synthesized by Microsynth AG. Two guides targeting one exon and the following intron of the respective gene locus of interest were designed for each KO cell line. Forward and reverse DNA oligonucleotides were annealed and inserted into the vector plasmid pX330-U6- Chimeric_BB_CBh_hSpCas9 (Addgene, 42230), using BbsI-HF® (NEB, R3539L) directed cloning. The sequence integrity was confirmed by Sanger sequencing. One day before transfection, 100 000 R1 mESCs per well were seeded in 12-well plates and the medium of the cells was replaced on the next day 1 h before transfection. Plasmid combinations, containing 2 sgRNAs (700 ng each) targeting one exon and one intron of each gene, were co-transfected with the fluorescent marker plasmid dsRed (100 ng) - as a proxy for transfection efficiency - using Lipofectamine™ 2000 Transfection Reagent (Invitrogen™, 11668019). The medium of transfected cells was changed after 6-12 h. Two to three days after transfection, single dsRed+ cells were FACS-sorted (BD FACSMelody™ Cell Sorter) onto a fibronectin-coated (10 µg/ml) 96-well plate to enable the generation of single clone KO cell lines. Successful KO of respective genes was confirmed by genotyping PCRs, mapping primers outside the deleted region to obtain a smaller fragment and combining outside-inside primers, resulting in a PCR product in the wild type but not in the KO. Genotyping was performed first with direct lysis reagent DirectPCR Lysis (Viagen Biotech, VIAG302-C) according to the data sheet to select clones for expansion. After clone selection and expansion based on genotyping, genomic DNA was extracted with the Puregene® Core Kit A (Qiagen, 158043) and genotyping PCRs were repeated and samples analyzed via Sanger sequencing (Supplementary Fig. 4). Additionally, genome editing was confirmed in immunofluorescence staining followed by imaging and Western blotting, if applicable.

### Generation of *Cep83/Cep83Δexon4* overexpression cell lines

For the generation of *Cep83/Cep83Δexon4* overexpression rescue cell lines, we cloned PiggyBac (pB) expression plasmids, containing *Cep83* or *Cep83Δexon4* RNA, isolated from wild type or *Cep83Δexon4* cell lines, respectively, under an EF1α promoter. Additionally, a N-terminal Flag-tag was added. Transfection of pB-Ef1α-Flag-Cep83-Ubi-Puro or pB-Ef1α-Flag-Cep83Δexon4-Ubi-Puro was utilized by Lipofectamine™ 2000 Transfection Reagent, using 500 ng of the construct and 1 µg PiggyBac transposase.

### Western Blot analysis

For Western blotting, cells were detached with trypsin from 6-well plates, the cell pellet washed with PBS and stored at −80°C after removal of the supernatant. Cells were lysed in 40 µl 1x RIPA buffer (Millipore, 20-188) containing cOmplete™ Protease Inhibitor Cocktail (Sigma-Aldrich, 11836145001) for 1 h on ice and vortexed every 10 min. Whole-cell extracts were collected by centrifugation (16,000 × g, 10 min, 4°C), cell debris removed and the protein concentration quantified using the Protein Assay Dye Kit (Bio-Rad, 500-0006). 30 µg of protein per sample was resolved on a 10% SDS-polyacrylamide gel and transferred to a PVDF membrane (Thermo Fisher Scientific, 88518), using the Wet/Tank Blotting System (Bio-Rad) at 400 mA for 1 h at 4°C. After Ponceau S staining of the membranes, they were washed in TBS (1x Tris-buffered saline with 0.1% Tween 20) and blocked with 5% milk in TBST for 30 min. Primary antibody incubation was performed overnight at 4°C, the secondary HRP conjugated antibodies incubated for 1 h at room temperature and the signal was detected with the Amersham™ ECL Select™ Western Blotting Detection Reagent (Cytiva, RPN2235).

### Propidium Iodide (PI) staining and FACS analysis

For cell cycle analysis, 150,000 cells were seeded in 6-well plates in 2iLIF in parallel to the mESC rosette cell assay. After 48 h, cells were detached with trypsin and counted. Cells were centrifuged for 5 min at 300 x g. Pellets with 500,000 cells per condition were first resuspended in 150 µl PBS to avoid clumping and 350 µl of 100% ice-cold ethanol was added dropwise while vortexing (final concentration 70% ethanol). Samples incubated for at least 30 min or overnight at 4°C and centrifuged for 5 min at 300 x g. For PI staining, pellets were resuspended in 300 µl PI-RNAse solution, containing 50 µg/ml PI (Sigma-Aldrich, P4170) and 100 µg/ml RNAse A (Thermo Fisher Scientific, EN0531), diluted in PBS and incubated for 20 min in the dark at room temperature. Single cell suspensions were obtained by transferring samples through a cell strainer cap (Corning) and measured with the LSRFortessa™ High-Parameter Flow Cytometer (BD Life Sciences - Biosciences). Data was analyzed using the FlowJoTM software (BD Life Sciences – Biosciences, version 10.5.3).

### Microscope specifications

Images were acquired with the following microscopes:

Visitron Live Spinning Disk, Spinning disc units: Yokogawa CSU-X1-A1 spinning disk (50 µm pinholes, spacing 253 µm, 5000 rpm), Yokogawa CSU-W1-T2 spinning disk (50 µm pinholes, spacing 500 µm, 4000 rpm), Cameras: EM-CCD (back-illuminated Andor iXon Life 888, 1024 x 1024 pixel, 13 µm pixel size, 16 bit, 26 fps (full frame), QE >95%) and sCMOS (back-illuminated Teledyne Prime BSI, 2048 x 2048 pixel, 6.5 µm pixel size, 16 bit, 43 fps (full frame), QE >95%), Objectives: CFI Plan Apo λ S 40xC/1.25 Sil, WD 0.30 mm (with coverslip thickness correction collar), CFI Plan Apo λ 60x/1.42 Oil, WD 0.15 mm, CFI Plan Apo λ 100x/1.45 Oil, WD 0.13 mm, Software VisiView 6.0,

Visitron Spinning Disk, Spinning disk unit: Yokogawa CSU-X1 Nipkow spinning disk unit (50 µm pinholes, spacing 253, 5000 rpm), Camera: sCMOS (70% QE, 2048 x 2048 pixel, 6.45 pixel size, 16 bit, up to 100 fps), Objective: Plan-Apochromat 63x/1.4 Oil DIC, WD 0.19 mm, Software VisiView 6.0

Zeiss LSM 980, Scanning mirrors with up to 8192 x 8192 pixels, Airyscan 2 compound detector consisting of 32 GaAsP detector units, Objective: Plan-Apochromat 63x/1.4 Oil DIC M27 (WD 0.19 mm), Software Zeiss ZEN 3.3

### Image processing and data analysis

Acquired images were analyzed using Fiji (Schindelin et al., 2012). Gastruloid growth and elongation was quantified based on morphology of images acquired daily using the ZOE Fluorescent Cell Imager (Bio-Rad). Perimeter, roundness (roundness = 4pi(area/perimeter^2)), Feret’s diameter and length were quantified in gastruloids by taking manual measurements with the polygon and segmented line tool (Supplementary Figure 2 A). Cilium length was determined by measuring the length of fluorescence for the ciliary marker ARL13B, ciliation rate (%) calculated based on counted cilia and DAPI staining to determine the cell number of the respective, quantified image. In gastruloids, the mean ratio of cilia to centrioles was determined based on immunofluorescence staining of gastruloids labeled with antibodies against ARL13B and γ-tubulin. This quantification method is not a direct readout of the ciliation rate per cell, due to difficulties in attributing centriole signal to individual cells in 3D samples, and was therefore only used to compare ciliogenesis in those samples. In 3D *in vitro* rosettes, we quantified lumen and rosette size based on masking the central z-plane (in-house developed Python code), using PODXL as a lumen marker and DAPI to assess the rosette size. Imaging conditions and subsequent post-acquisition processing was always constant within each experiment for image analysis. For representative visual depiction in Figures, images were cropped, brightness and contrast optimized using Fiji.

### Graphing and statistics

All graphs were created using GraphPad Prism 10 and statistically analyzed using a two-tailed unpaired Student’s t-test (normal distributed data) for independent data sets including at least 3 experimental replicates. P values: ns > 0.05, *p ≤ 0.05, **p ≤ 0.01, ***p ≤ 0.001, ****p ≤ 0.0001.

## Supporting information

Supplemental Informatiom

## Acknowledgements

We would like to thank all members of the Buecker lab for discussions and feedback through the project, including the summer student Laura Rüland for her help, the Dammermann and Leeb labs for shared lab meetings/journal clubs, critical input and valuable support, the BioOptics-FACS and BioOptics-Light Microscopy facility at Max Perutz Labs, Nicholas Wedige and Lorenz Perschy for help with the image analysis pipeline of 3D *in vitro* rosettes. This work was supported by the Austrian Science Fund FWF (PAT9017923, P34123, W1261 and DOC72 to CB and F8803-B to AD).

## Author contributions

C.B. and I.V. conceived and designed the study, with critical input and co-supervision from A.D.; I.V. carried out and analyzed all experiments unless otherwise specified in this paragraph; T.C. performed the majority of gastruloid experiments including image analysis and quantifications, cDNA sequencing of *Cep83* mutants, generated pB-transfected Flag-*Cep83Δexon4* and *Flag-Cep83* overexpression cell lines and executed all experiments with these cell lines, under supervision of I.V.; M.E. carried out starvation experiments of wild type cells and p27 Western blots under supervision of I.V.; M.H. and S.F. generated the *Cep83^-/-^*cell line and contributed to genotyping; I.V. wrote the manuscript and designed all figures, with input from all members of the Buecker lab and A.D.; C.B. supervised the project.

## Supplementary Figure Legends

**S1 Fig. Gastruloids express SOX2, Brachyury and SOX17-positive cells, related to Figure 5**

(A) Spatiotemporal expression of gastruloid markers. Representative example of immunofluorescence staining of gastruloids at 72, 96 and 120 h, labeled with antibodies against SOX2 (red), Brachyury (green) and DAPI staining (blue). Maximum intensity projection is depicted. (B) Representative example of immunofluorescence staining of gastruloids at 120 h, labeled with antibodies against SOX17 (green) and DAPI staining (blue). The central z-plane is depicted. Scale bar (A, B): 100 µm.

**S2 Fig. Schematic of measurements and length of *Ift88* KO gastruloids, related to Figure 6**

(A) Morphology-based quantification of gastruloids: The dashed line (yellow) was used to quantify perimeter and roundness (roundness = 4pi(area/perimeter^2)) of gastruloids. Feret’s diameter (red line) and the major axis length (green) indicate gastruloid length. Roundness: Roundness = 4*area/(π*major_axis^2). Perfect circles = score of 1 (due to measurement by polygon tool, perfect circles achieve a score of 0,995). Length: diameter up to 96 h gastruloids and longest measured middle axis for 120 h gastruloids. (B) Box plots indicating the length of WT and *Ift88* KO gastruloids. The box represents the interquartile range (IQR), with the median indicated by a horizontal line. Whiskers extend to 1.5 × IQR, and data points beyond this range are considered outliers (shown as dots). n=3 independent experiments with one representative experiment depicted.

**S3 Fig. *Cep83* is not essential for gastruloid formation, related to Figure 7**

(A, B) Box plots indicating the length of WT, *Cep83^-/-^*and *Cep83Δexon4* gastruloids. The box represents the interquartile range (IQR), with the median indicated by a horizontal line. Whiskers extend to 1.5 × IQR, and data points beyond this range are considered outliers (shown as dots). n=3 independent experiments with one representative experiment depicted. (C, D) Immunofluorescence staining of WT, *Cep83^-/-^* and *Cep83Δexon* gastruloids at 72, 96 and 120 h, labeled with antibodies against SOX2 (red), Brachyury (green) and DAPI staining (blue). The maximum intensity projection is depicted. Scale bar (C, D): 100 µm.

**S4 Fig. Sanger sequencing of *Plk4* KO, *Ift88* KO and *Cep83^-/-^* cell lines indicates CRISPR/Cas9-mediated deletions**

Sanger sequencing of gDNA: (A) *Plk4* KO clone 1 and 2 show 844 bp deletion at the gRNA target site, (B) *Ift88* KO clone 1 shows 866 bp deletion and 9 bp insertion, *Ift88* KO clone 2 shows 799 bp deletion, (C) *Cep83^-/-^* clone 1 and 2 show 5983 bp deletion, (D) *Cep83Δexon4* clone 4 shows 857 bp deletion.

**S5 Fig. *Cep83Δexon4* overexpression does not restore ciliogenesis in 2D mESCs and partially rescues ciliation in gastruloids, related to Figure 8**

(A) Representative example of immunofluorescence staining of WT, *Cep83Δexon4*, *Cep83Δexon4 with Cep83 overexpression*, *Cep83Δexon4* with *Cep83Δexon4* overexpression after induced ciliogenesis (24 h of starvation), labeled with antibodies against γ-tubulin (red), ARL13B (green) and DAPI staining (blue). Maximum intensity projection depicted. (B) Quantification of the mean ratio of cilia to centrioles in WT, *Cep83Δexon4*, WT with *Cep83* overexpression, *Cep83Δexon4* with *Cep83* overexpression, *Cep83Δexon4* with *Cep83Δexon4* overexpression after 72 h of gastruloid formation. Bars represent the mean ratio between replicates. n = 3 independent experiments; note that for the WT + *Cep83* OE, *Cep83Δexon4* + *Cep83* OE, and *Cep83Δexon4* + *Cep83Δexon4* OE conditions there is only one replicate. (C) Sequenced cDNA of *Cep83Δexon4* cells shows that in all clones, exon 4 is skipped. Clone 3 is shown representatively. Scale bar (A): 20 µm.

## Notes

### Competing Interest Statement

The authors have declared no competing interest.

