## Supplemental Informatiom for "*In vitro* approaches to study centriole and cilium function in early mouse embryogenesis"

S1 Fig.

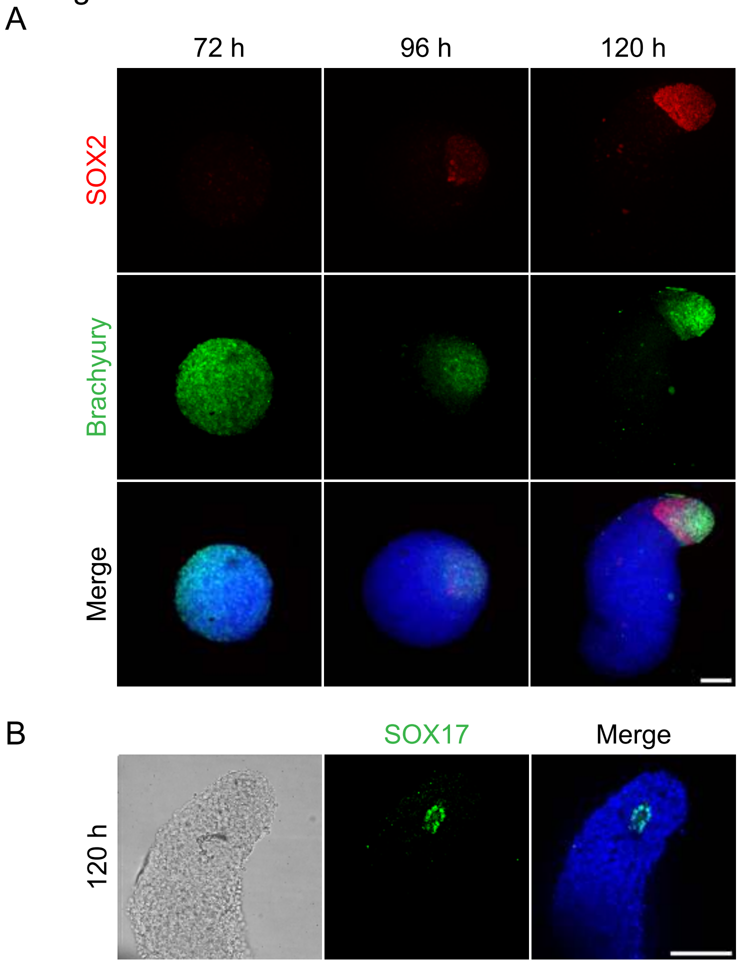

S2 Fig.

A

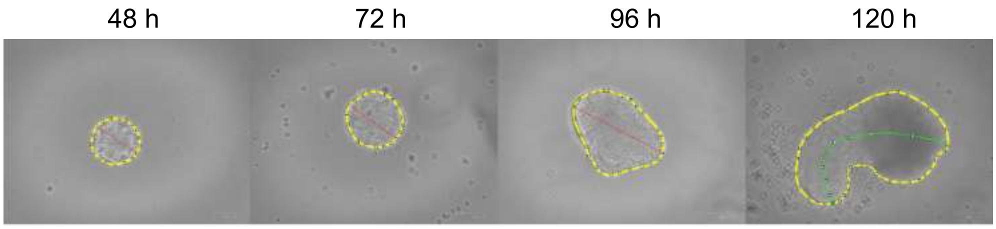
48 h
72 h
96 h
120 h

B

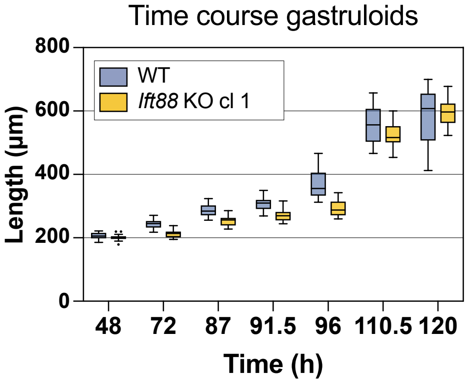
Time course gastruloids
800
600
400
200
0
Length (µm)
WT
Ift88 KO cl 1
48
72
87
91.5
96
110.5
120
Time (h)

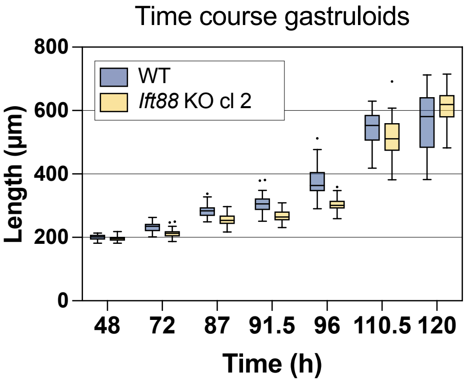
Time course gastruloids
800
600
400
200
0
Length (µm)
WT
Ift88 KO cl 2
48
72
87
91.5
96
110.5
120
Time (h)

S3 Fig.

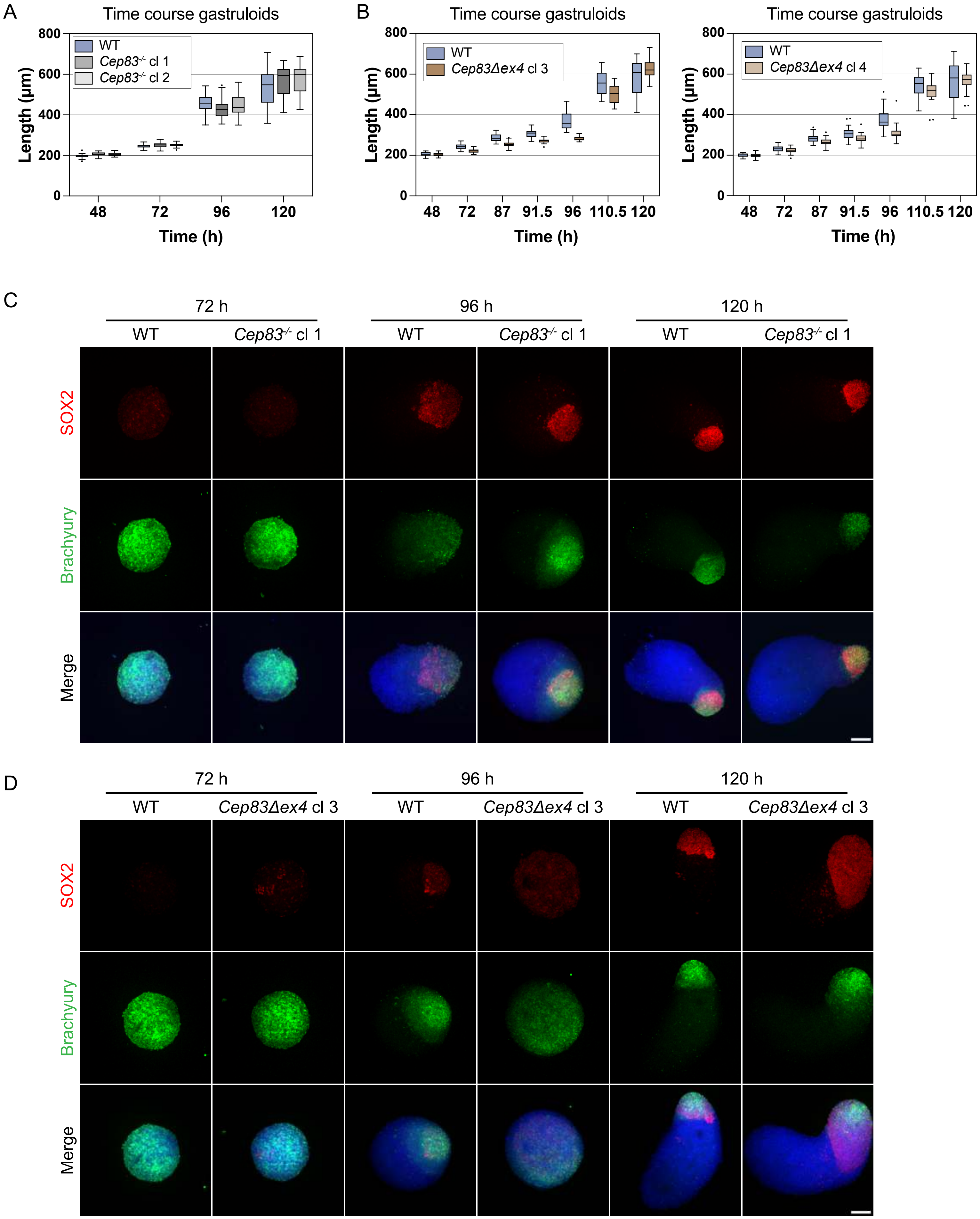
A
Time course gastruloids
WT
Cep83-/- cl 1
Cep83-/- cl 2
Length (µm)
0
200
400
600
800
48
72
96
120
Time (h)
B
Time course gastruloids
WT
Cep83Δex4 cl 3
Length (µm)
48
72
87
91.5
96
110.5
120
Time (h)
Time course gastruloids
WT
Cep83Δex4 cl 4
Length (µm)
Time (h)
C
72 h
96 h
120 h
WT
Cep83-/- cl 1
SOX2
Brachyury
Merge
D
72 h
96 h
120 h
WT
Cep83Δex4 cl 3
SOX2
Brachyury
Merge

S4 Fig.

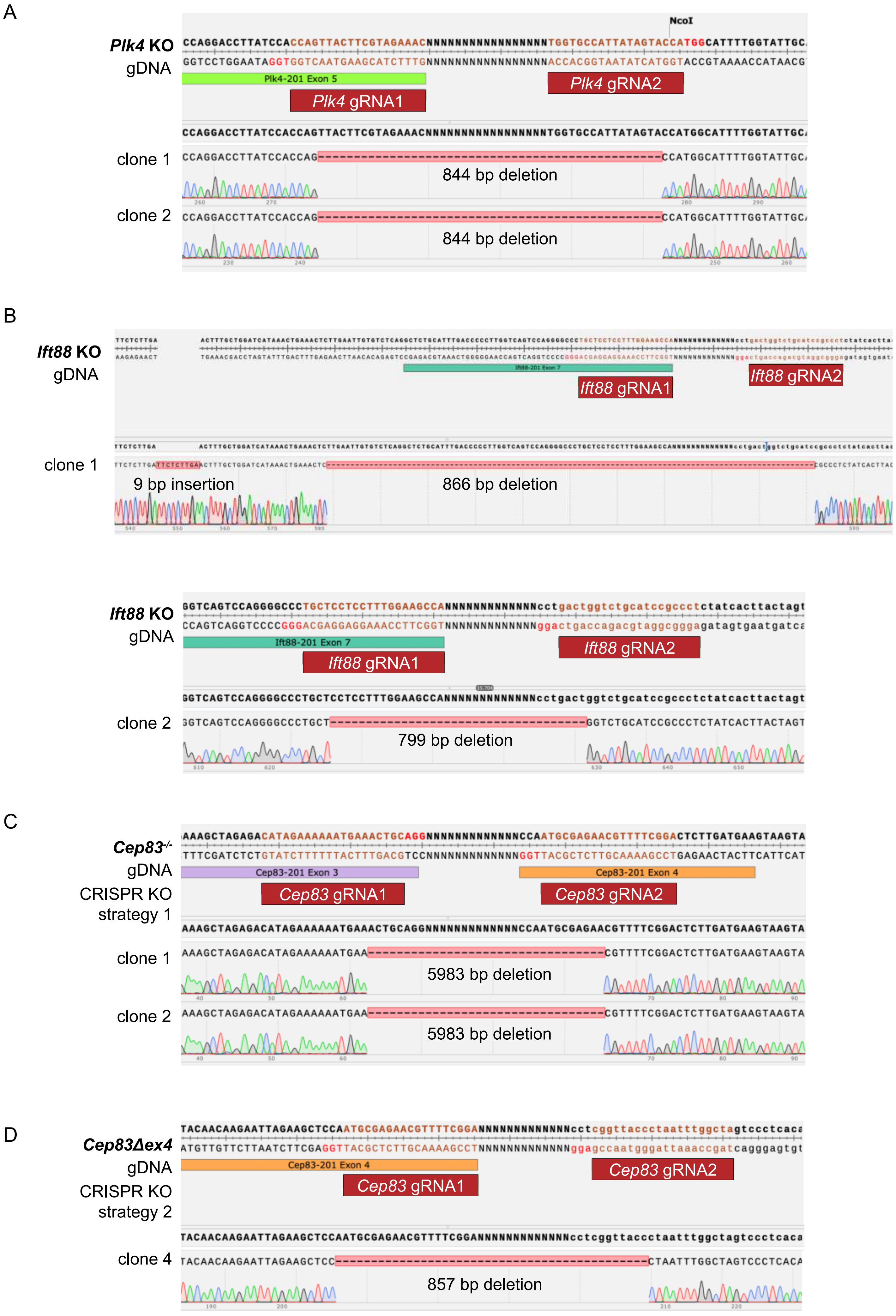
A
NcoI
Plk4 KO
gDNA
Plk4-201 Exon 5
Plk4 gRNA1
Plk4 gRNA2
clone 1
844 bp deletion
clone 2
844 bp deletion
B
Ift88 KO
gDNA
Ift88 gRNA1
Ift88 gRNA2
clone 1
9 bp insertion
866 bp deletion
Ift88 KO
gDNA
Ift88-201 Exon 7
Ift88 gRNA1
Ift88 gRNA2
clone 2
799 bp deletion
C
Cep83-/-
gDNA
CRISPR KO
strategy 1
Cep83-201 Exon 3
Cep83-201 Exon 4
Cep83 gRNA1
Cep83 gRNA2
clone 1
5983 bp deletion
clone 2
5983 bp deletion
D
Cep83Δex4
gDNA
CRISPR KO
strategy 2
Cep83-201 Exon 4
Cep83 gRNA1
Cep83 gRNA2
clone 4
857 bp deletion

S5 Fig.

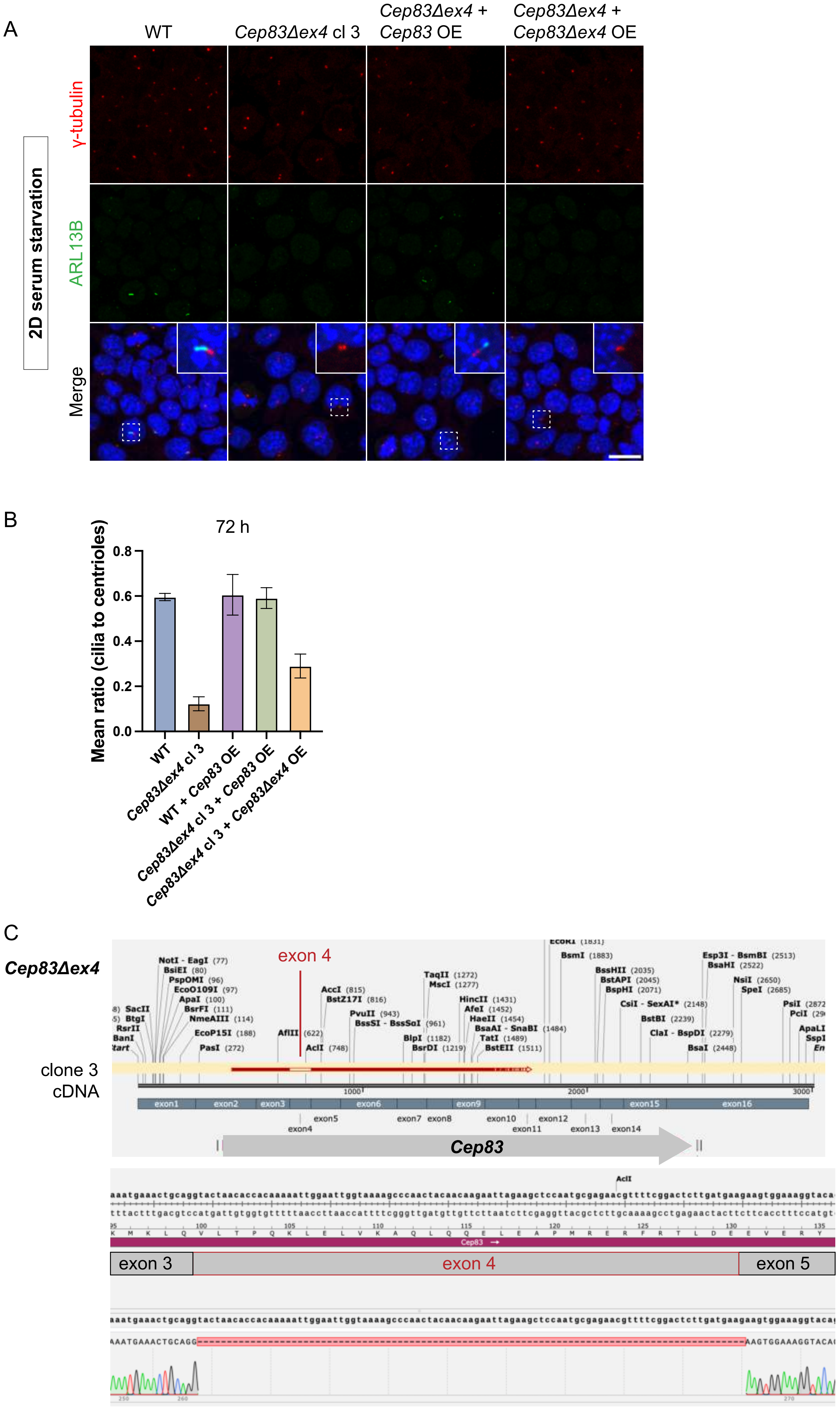

A
WT
Cep83Δex4 cl 3
Cep83Δex4 + Cep83 OE
Cep83Δex4 + Cep83Δex4 OE
2D serum starvation
γ-tubulin
ARL13B
Merge
B
72 h
Mean ratio (cilia to centrioles)
0.0
0.2
0.4
0.6
0.8
WT
Cep83Δex4 cl 3
WT + Cep83 OE
Cep83Δex4 cl 3 + Cep83 OE
Cep83Δex4 cl 3 + Cep83Δex4 OE
C
Cep83Δex4
clone 3 cDNA
exon 4
NotI - EagI (77)
BsiEI (80)
PspOMI (96)
EcoO109I (97)
ApaI (100)
BsrFI (111)
NmeAIII (114)
EcoP15I (188)
PasI (272)
AflII (622)
AclI (748)
AccI (815)
BstZ17I (816)
PvuII (943)
BssSI - BssSoI (961)
BlpI (1182)
BsrDI (1219)
TaqII (1272)
MscI (1277)
HincII (1431)
AfeI (1452)
HaeII (1454)
BsaAI - SnaBI (1484)
TatI (1489)
BstEII (1511)
EcoRI (1831)
BsmI (1883)
BssHII (2035)
BstAPI (2045)
BspHI (2071)
CsiI - SexAI* (2148)
BstBI (2239)
ClaI - BspDI (2279)
BsaI (2448)
Esp3I - BsmBI (2513)
BsaHI (2522)
NsiI (2650)
SpeI (2685)
1000
2000
3000
exon1
exon2
exon3
exon4
exon5
exon6
exon7
exon8
exon9
exon10
exon11
exon12
exon13
exon14
exon15
exon16
Cep83
AclI
aaatgaaactgcaggtactaacaccacaaaaattggaattggtaaaagcccaactacaacaagaattagaagctccaatgcgagaacgttttcggactcttgatgaagaagtggaaaggtaca
tttactttgacgtccatgattgtggtgtttttaaccttaaccattttcgggttgatgttgttcttaatcttcgaggttacgctcttgcaaaagcctgagaactacttcttcacctttccatgt
100
105
110
115
120
125
130
135
K M K L Q V L T P Q K L E L V K A Q L Q E L E A P M R E R F R T L D E E V E R Y
Cep83
exon 3
exon 4
exon 5
aaatgaaactgcaggtactaacaccacaaaaattggaattggtaaaagcccaactacaacaagaattagaagctccaatgcgagaacgttttcggactcttgatgaagaagtggaaaggtaca
AAATGAAACTGCAGG
AAGTGGAAAGGTACA
250
260
270

| Reagent/Resource | Source | Identifier (Cat#) |
|---|---|---|
| **Chemicals, enzymes and reagents** | | |
| AlbuMAX™ II | Gibco™ | 11021029 |
| Amersham™ ECL Select™ Western Blotting Detection Reagent | Cytiva | RPN2235 |
| BbsI-HF® | NEB | R3539L |
| B-27™ Supplement | Gibco™ | 17504044 |
| CHIR99021 | MedChemExpress | HY-10182 |
| cOmplete™ Protease Inhibitor Cocktail | Sigma-Aldrich | 11836145001 |
| Cultrex Reduced Growth Factor Basement Membrane Extract, Type 2 | R&D Systems | 3533-005-02 |
| DAPI | Sigma-Aldrich | D9542 |
| DirectPCR Lysis reagent | Viagen Biotech | VIAG302-C |
| DMEM/F-12, with GlutaMAX™ | Gibco™ | 31331028 |
| Doxycycline Hyclate | Sigma-Aldrich | D9891 |
| Dulbecco's Phosphate-Buffered Saline | Sigma-Aldrich | D8537 |
| Fetal Bovine Serum | Sigma-Aldrich | F7524 |
| Fibronectin | YO Proteins AB | 663 |
| GlutaMAX™ Supplement, 200mM (=L-Alanyl-Glutamine) | Gibco™ | 3133102 |
| hLIF | provided by the VBCF Protein Technologies Facility | https://www.viennabiocenter.org/facilities/ |
| HyClone™ DMEM/F12 1:1 mix | Cytiva | SH30271.FS |
| Laminin | Sigma-Aldrich | L2020 |
| Lipofectamine™ 2000 Transfection Reagent | Invitrogen™ | 11668019 |
| MycoAlertTM PLUS Mycoplasma Detection Kit | Lonza | LT07-705 |
| MEM NEAA | Gibco™ | 11140035 |
| Neurobasal™ Medium | Thermo Fisher Scientific | 21103049 |
| N-2 Supplement, 100X | Gibco™ | 17502048 |
| Paraformaldehyde 16% aq soln, methanol free | MP Biomedicals | 219998380 |
| PD0325901 | MedChemExpress | HY-10254 |
| Penicillin-Streptomycin | Gibco™ | 15070063 |
| Poly-L-ornithine hydrobromide | Sigma-Aldrich | P4638 |
| Propidium Iodide (PI) | Sigma-Aldrich | P4170 |
| Puregene® Core Kit A | Qiagen | 158043 |
| PVDF membrane | Thermo Fisher Scientific | 88518 |
| RIPA buffer | Millipore | 20-188 |
| RNase A | Thermo Fisher Scientific | EN0531 |

| | | |
|---|---|---|
| Sodium Pyruvate | Gibco™ | 11360039 |
| Triton™ X-100 | PanReac AppliChem ITW Reagents | 142314 1611 |
| Trypsin-EDTA solution | Sigma-Aldrich | T3924 |
| TWEEN® 20 | Sigma-Aldrich | P7949 |
| VECTASHIELD® Antifade Mounting Medium with DAPI | Vector Laboratories | VEC-H-1200 |
| 2-Mercaptoethanol | Gibco™ | 21985023 |

| **Equipment** | | |
|---|---|---|
| BD FACSMelody™ Cell Sorter | | N/A |
| ChemiDoc™ Touch Imaging System | Bio-Rad | N/A |
| Corning™ 96-Well Clear Ultra Low Attachment Microplates | Corning™ | 7007 |
| CytoOne® Multi-Well Plates 6-well | CytoOne | CC7682-7506 |
| CytoOne® Multi-Well Plates 12-well | CytoOne | CC7682-7512 |
| CytoOne® Multi-Well Plates 96-well | CytoOne | CC7682-7596 |
| LSRFortessa™ High-Parameter Flow Cytometer | BD Life Sciences - Biosciences | N/A |
| Multi-Well Plates 24-well | SARSTEDT | 833.922 |
| Multi-Well Plates 48-well | SARSTEDT | 833.923 |
| µ-Slide 8 Well Glass Bottom Plate | ibidi | 80827 |
| ZOE Fluorescent Cell Imager | Bio-Rad | N/A |

| **Experimental models: Cell lines** | | |
|---|---|---|
| R1 mouse embryonic stem cells (WT) | Nagy et al., 1993 | N/A |
| *Ift88* KO clone 1 | This study | N/A |
| *Ift88* KO clone 2 | This study | N/A |
| *Plk4* KO clone 1 | This study | N/A |
| *Plk4* KO clone 2 | This study | N/A |
| *Cep83$^{-/-}$* KO clone 1 | This study | N/A |
| *Cep83$^{-/-}$* KO clone 2 | This study | N/A |
| *Cep83Δexon4* clone 3 | This study | N/A |
| *Cep83Δexon4* clone 4 | This study | N/A |
| *Cep83Δexon4 Cep83 OE* clone 3 | This study | N/A |
| *Cep83Δexon4 Cep83Δexon4* clone 3 | This study | N/A |
| WT *Cep83* OE | This study | N/A |

| **Recombinant DNA** | | |
|---|---|---|
| pX330-U6-Chimeric_BB_CBh_hSpCas9 | Chong et al., 2013 | Addgene Plasmid #42230 |
| pB Ef1 Flag-CEP83 Ubi Puro | This study | N/A |

| | | |
|---|---|---|
| pB Ef1 Flag-CEP83deltaExon4 Ubi Puro | This study | N/A |

| gRNA sequences | Sequence |
|---|---|
| Ift88 gRNA 1 forward | CACCGTGGCTTCCAAAGGAGGAGCA |
| Ift88 gRNA 1 reverse | AAACTGCTCCTCCTTTGGAAGCCAC |
| Ift88 gRNA 2 forward | CACCGAGGGCGGATGCAGACCAGTC |
| Ift88 gRNA 2 reverse | AAACGACTGGTCTGCATCCGCCCTC |
| Plk4 gRNA 1 forward | CACCGGTTTCTACGAAGTAACTGG |
| Plk4 gRNA 1 reverse | AAACCCAGTTACTTCGTAGAAACC |
| Plk4 gRNA 2 forward | CACCGTGGTGCCATTATAGTACCA |
| Plk4 gRNA 2 reverse | AAACTGGTACTATAATGGCACCAC |
| Cep83 strategy 1 gRNA 1 forward | CACCGTCCGAAAACGTTCTCGCAT |
| Cep83 strategy 1 gRNA 1 reverse | AAACATGCGAGAACGTTTTCGGAC |
| Cep83 strategy 1 gRNA 2 forward | CACCGCATAGAAAAAATGAAACTGC |
| Cep83 strategy 1 gRNA 2 reverse | AAACGCAGTTTCATTTTTTCTATGC |
| Cep83 strategy 2 gRNA 1 forward | CACCGTCCGAAAACGTTCTCGCAT |
| Cep83 strategy 2 gRNA 1 reverse | AAACATGCGAGAACGTTTTCGGAC |
| Cep83 strategy 2 gRNA 2 forward | CACCGTAGCCAAATTAGGGTAACCG |
| Cep83 strategy 2 gRNA 2 reverse | AAACCGGTTACCCTAATTTGGCTAC |

| Primers | Sequence |
|---|---|
| *Ift88* KO validation 1 primer 1 forward | ACCCCAGCACTTTCGAGGCT |
| *Ift88* KO validation 1 primer 2 reverse | TGTGCTGCCCTAACAACTCCACT |
| *Ift88* KO validation 1 primer 3 reverse | ATGCGTGCGTGTTTGAGGCA |
| *Plk4* KO validation 1 primer 1 forward | TTGCACCGGGACCTCACACT |
| *Plk4* KO validation 1 primer 2 reverse | AAGGAAGGGTTGGGGACAGAACC |
| *Plk4* KO validation 1 primer 3 forward | ACCAGTTTGAGTGGCAGCCTACTT |
| *Cep83*$^{-/-}$ KO validation 1 primer 1 forward | GGTCTGGAGGTCGCTGATTA |
| *Cep83*$^{-/-}$ KO validation 1 primer 2 reverse | TGGAAGCAGTCAGAGTAGGC |
| *Cep83*$^{-/-}$ KO validation 2 primer 3 forward | ACTAGAGTTGTGTGGGCTGC |
| *Cep83*$^{-/-}$ KO validation 2 primer 4 reverse | GGGTGGAGAAATGGCATTGG |
| *Cep83*$^{-/-}$ KO validation 3 primer 5 forward | TGAAAAGCAAGCAAGCCAGG |
| *Cep83*$^{-/-}$ KO validation 3 primer 6 reverse | GTCCTGTCCCATTATGCCTCAG |
| *Cep83*$^{-/-}$ KO validation 4 primer 7 forward | GGTCTGGAGGTCGCTGATTA |
| *Cep83*$^{-/-}$ KO validation 4 primer 8 reverse | GGGTGGAGAAATGGCATTGG |
| *Cep83*$^{-/-}$ KO validation 5 primer 9 forward | ACTAGAGTTGTGTGGGCTGC |
| *Cep83*$^{-/-}$ KO validation 5 primer 10 reverse | TGGAAGCAGTCAGAGTAGGC |
| *Cep83*$^{-/-}$ KO validation 6 primer 11 forward | GCTGCGGCTTTGCTCTGGAA |
| *Cep83*$^{-/-}$ KO validation 6 primer 12 reverse | TCTGGTGGGTGGTGGGTTGAAA |
| *Cep83Δexon4* validation 1 primer 1 forward | GCATCCATGGCGTGTGTGACA |
| *Cep83Δexon4* validation 1 primer 2 reverse | AGCAGCAGAGCAGTCCTGTGT |

| | |
|---|---|
| *Cep83Δexon4* validation 1 primer 3 reverse | ATCACGGTGGAGACGGCACA |
| Age1_Kozak_Start_Flag _Linker_CEP83 Fw | GGGACCGGTGCCACCATGGACTACA AGGACGACGATGACAAGGGCGGAG GTGGTTCTGGCGGTGGAGGTTCGGA CACATTTCCTAGCCTATTTCCAC |
| Not1_Xho1_Cep83_rev | AAATCGCGGCCGCTCGAGTCACTCCC CAGGAGACCC |

| Reagent/Resource | Dilution | Source | Identifier (Cat#) |
|---|---|---|---|
| **Primary antibodies (IF staining)** | | | |
| Anti-ARL13B Rabbit pAb | 1:500 | Proteintech | 17711-1-AP |
| Anti-Acetylated-α-Tubulin Mouse mAb | 1:500 | Sigma-Aldrich | T6793 |
| Anti-Brachyury Goat pAb | 1:100 | R&D Systems | AF2085 |
| Anti-Podocalyxin (Podxl) Rat mAb | 1:500, 1:300 (gastruloids) | R&D Systems | MAB1556 |
| Anti-Sox2 Rat mAb | 1:300 | eBioscience™ | 15218187 |
| Anti-Sox17 Goat pAb | 1:100 | R&D Systems | AF1924 SP |
| Anti-γ-Tubulin Mouse mAb | 1:300, 1:150 (gastruloids) | Sigma-Aldrich | T6557 |

| Reagent/Resource | Dilution | Source | Identifier (Cat#) |
|---|---|---|---|
| **Secondary antibodies** | | | |
| Anti-Goat IgG H&L (Alexa Fluor® 488) Donkey | 1:500 | abcam | ab150129 |
| Anti-Mouse IgG H&L (Alexa Fluor® 555) Donkey | 1:500 | abcam | ab150106 |
| Anti-Mouse IgG H&L (Alexa Fluor® 647) Donkey | 1:500 | abcam | ab150107 |
| Anti-Rabbit IgG H&L (Alexa Fluor® 488) Donkey | 1:500 | abcam | ab150073 |
| Anti-Rat IgG H&L (Alexa Fluor® 555) Donkey | 1:500 | abcam | ab150154 |
| Anti Rabbit IgG H&L (Alexa Fluor® 488) Goat | 1:500 | abcam | ab150077 |
| Anti-Mouse IgG H&L (Alexa Fluor® 647) Goat | 1:500 | abcam | ab150115 |

| Reagent/Resource | Dilution | Source | Identifier (Cat#) |
|---|---|---|---|
| **Antibodies (Western blot)** | | | |
| Anti-CEP83 Rabbit pAb | 1:1000 | Proteintech | 26013-1-AP |
| Anti-p27 KIP1 Rabbit mAb | 1:1000 | abcam | ab32034 |
| Anti-Vinculin Mouse mAb | 1:5000 | Santa Cruz | sc-73614 |
| Anti-Mouse IgG H&L (HRP) Goat | 1:10000 | abcam | ab205719 |
| Anti-Rabbit IgG H&L (HRP) Goat | 1:10000 | abcam | ab6721 |

| Software | | |
|---|---|---|
| FlowJo™ | BD Life Sciences – Biosciences | https://www.flowjo.com/solutions/flowjo version 10.5.3 |
| GraphPad Prism | Dotmatics | https://www.graphpad.com/features |
| Fiji | Schindelin et al., 2012 | https://fiji.sc |
